# Spectral graph theory efficiently characterises ventilation heterogeneity in lung airway networks

**DOI:** 10.1101/2020.04.15.042416

**Authors:** Carl A. Whitfield, Peter Latimer, Alex Horsley, Jim M. Wild, Guilhem J. Collier, Oliver E. Jensen

## Abstract

This paper introduces a linear operator for the purposes of quantifying the spectral properties of transport within resistive trees, such as airflow in lung airway networks. The operator, which we call the Maury matrix, acts only on the terminal nodes of the tree and is equivalent to the adjacency matrix of a complete graph summarising the relationships between all pairs of terminal nodes. We show that the eigenmodes of the Maury operator have a direct physical interpretation as the relaxation, or resistive, modes of the network. We apply these findings to both idealised and image-based models of ventilation in lung airway trees and show that the spectral properties of the Maury matrix characterise the flow asymmetry in these networks more concisely than the Laplacian modes, and that eigenvector centrality in the Maury spectrum is closely related to the phenomenon of ventilation heterogeneity caused by airway narrowing or obstruction. This method has applications in dimensionality reduction in simulations of lung mechanics, as well as for characterisation of models of the airway tree derived from medical images.

## 1 Background

In healthy human lungs, the airways form a bifurcating tree where, on average, around the first 16 generations of airways are purely conductive and serve to transport gas from the mouth to the alveolar region where the majority of gas exchange takes place. The conducting airways terminate in ∼30,000 respiratory units (or acini) [1], where diffusion becomes the dominant transport mechanism. Resistive flow in the conducting airways, as well as tissue compliance, determine the ventilation mechanics and hence require a high-dimensional mathematical model to be represented accurately [2, 3, 4]. Efficient dimensionality reduction algorithms are needed to improve the computation time and tractability of using such models in inverse problems in the future.

Furthermore, understanding the relationship between structure and function in the lungs is crucial to identifying lung disease physiology and its progression. One particular marker of progression in obstructive lung conditions is “ventilation heterogeneity” (VH), or the uneven distribution of fresh gas in the lungs, which can result from obstructions in the airways [5, 6, 7]. The methods presented here provide a new approach to characterise the resistance structure and its effect on VH. This will be useful for visualisation and characterisation of complex airway models based on patient computed tomography (CT) images, lung casts or micro-CT scans of excised lungs.

Graph theoretic perspectives have proven useful in characterising, classifying and understanding transport in various examples of biological networks [8, 9, 10]. The approach we take in this paper uses concepts from spectral graph theory, where a linear operator describing a particular physical property or characteristic of the network is decomposed into its eigenspectum. This has proved useful for dimensionality reduction and visualisation of large data sets in various areas of science [11] and has a number of real-world applications [12] including diffusive transport networks [13], but has not previously been applied to the airways. Some standard operators, such as the adjacency and Laplacian operators, are generic to all networks and can be adapted to characterise different processes or uncover important motifs in the network [14]. However, for lung airway networks and other resistive trees, these operators are not necessarily the optimal representation of the physical processes of interest on the network, as we show here.

In this paper we demonstrate that the resistive properties of realistic tree networks are characterised by a tree-specific operator that we call the Maury matrix, first introduced in ref. [15]. This matrix provides a complete description of the linear resistance relations on the network. In section 2, we formulate the flow problem on an airway network in terms of the conductance Laplacian and the Maury matrix, and show how each can be approximated via spectral reduction. We show how VH is evaluated for a set of realistic airway network models (four of which are based on CT imaging). In section 3, we compare spectral approximations of the Laplacian and Maury operators, and show in particular how the reduced Maury operator efficiently captures spatial patterns of VH, giving a new method for dimensionality reduction in these systems.

## 2 Methods

Unlike several other examples of biological transport networks, mammalian airways develop robustly into tree networks [1], such that they contain no cycles and airways can be defined hierarchically by parent airways branching into children airways. Therefore, we model a tree network 𝒩 = {𝒱, ℰ} consisting of a set of nodes (or vertices), 𝒱 = {*v*_*i*_}, and a set of branches (or edges) representing airways, ℰ = {*e*_*j*_ = (*v*_*i*_, *v*_*i*′_)}. We define each branch’s orientation to point from its more proximal node *v*_*i*_ to the more distal node *v*_*i*′_ based on its graph distance from the root node *v*_1_.

The set of terminal nodes is denoted 𝒯 ⊂ 𝒱 and we order the indices *i* so that the terminal nodes are listed last 𝒯 = {*v*_|𝒱|−|𝒯 |+1_, …, *v*_|𝒱|_}, where |.| indicates the number of objects in the set. The tree topology means that every branch *e*_*j*_ has a unique distal node *v*_*i*′_ so we index the branches such that *j* = *i*′ − 1. In the conducting airways of the lung, the root node represents the entrance from the upper airway, which is connected to the trachea that we label as branch *e*_1_. See figure 1(a) for an overview of this notation. The distribution of flow through the airway network in the lung is primarily dependent on the interplay between resistance and compliance of the airways and tissue. In this paper we focus on new ways to model and analyse the resistance of the airway network, and its implications for the distribution of gas flow.

**Figure 1:**
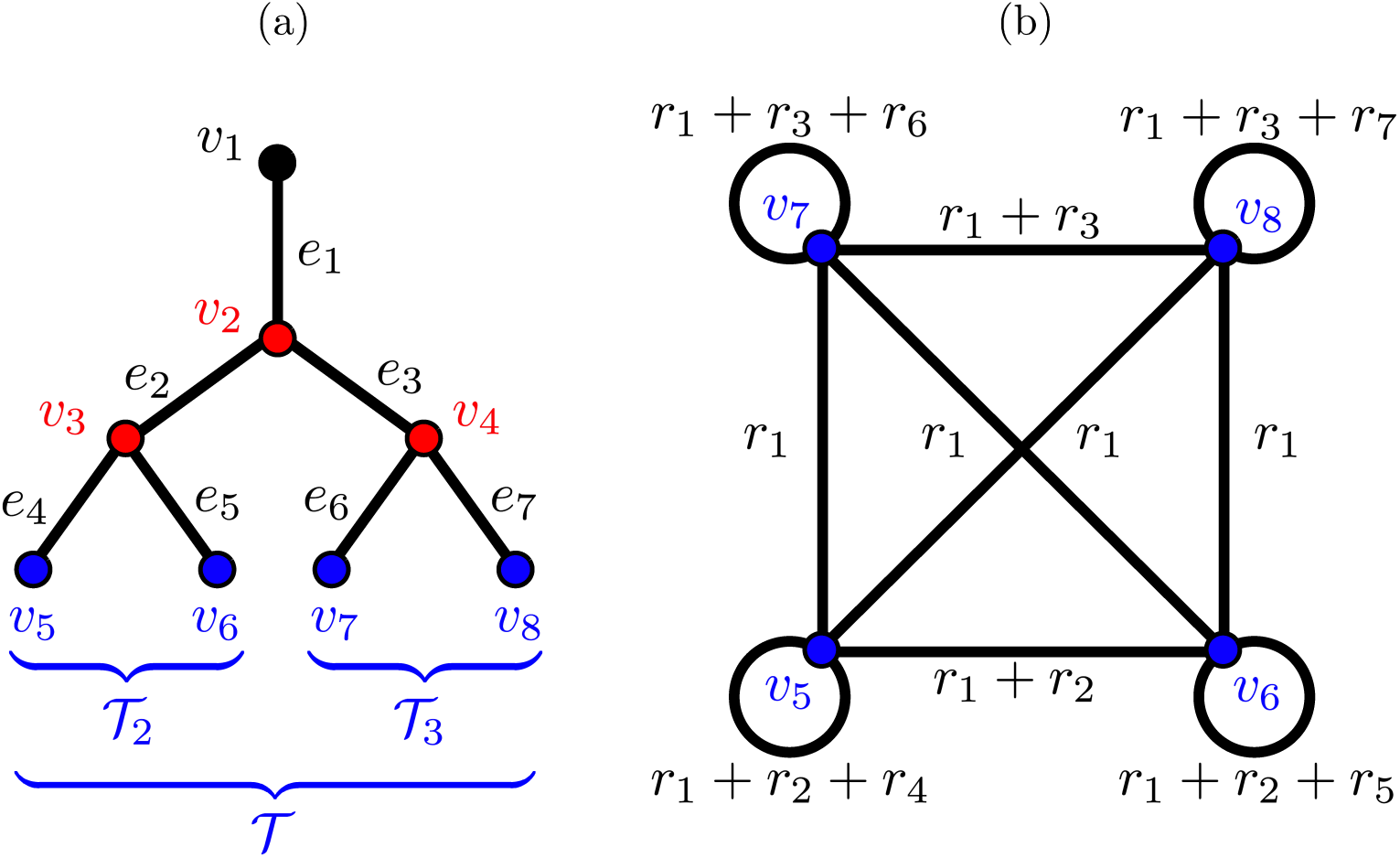
(a) Sketch of an airway tree network with nodes 𝒱 = {*v*_1_, …, *v*_8_} and branches ℰ = {*e*_1_, …, *e*_7_}. The set of internal nodes 𝒱_int_ = {*v*_2_, *v*_3_, *v*_4_} is coloured red and the set of terminal nodes 𝒯 = {*v*_5_, …, *v*_8_} is coloured blue. The sets 𝒯_2_ and 𝒯_3_ denote the subsets of 𝒯 descended from branchs *e*_2_ and *e*_3_ respectively. (b) The network shown here has weighted adjacency matrix **R** where the nodes are the terminal nodes 𝒯 of the airway tree. The branch weights are labelled where *r*_*j*_ corresponds to the resistance of branch *e*_*j*_ in the airway tree.

### 2.1 The Conductance Laplacian

The network incidence matrix **N** ∈ ℤ^|𝒱|*×*|ℰ|^ maps the nodes of the network to their associated branches. Its entries are

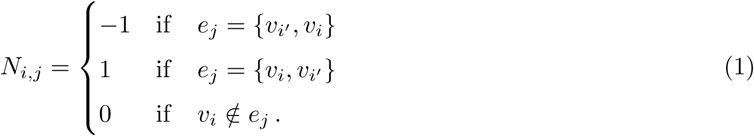

We model flow through the conducting airways of the lung by the linear resistance equation

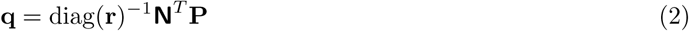

where **q** ∈ ℝ^|ℰ|^ is the vector of branch fluxes, **r** ∈ ℝ^| ℰ|^ is the vector of branch resistances and **P** ∈ ℝ^| 𝒱|^ is the vector of node pressures. The incidence matrix **N** is a discrete boundary operator, mapping 1-chains (branch quantities) to 0-chains (node quantities), and is the discrete analogue of the divergence operator in vector calculus [16]. Similarly, **N**^*T*^ is analogous to the gradient operator and performs the reverse mapping. Applying **N** to (2) gives

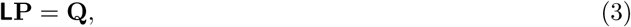

where **L** = **N**diag(**r**)^−1^**N**^*T*^ is the network conductance Laplacian. The node flux **Q** = **Nq** for each node *v*_*i*_ is equal to the sum of fluxes into node *i* from the branches connected to it. Assuming incompressibility, these are zero except on any nodes that are sources or sinks. In the lung, gas can only enter or exit the conducting airways via the upper airway or the transitional bronchioles (which feed the acini), so henceforth, we assume that the root *v*_1_ and terminal nodes *v*_*i*_ ∈ 𝒯 are the only sources or sinks. Therefore, the vectors of node and branch quantities can be written as

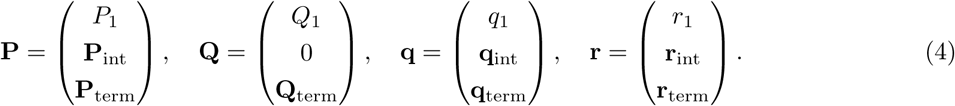

Here, the separate blocks of **P** and **Q** distinguish the quantities defined on the root node *v*_1_, internal nodes 𝒱_int_ = {*v*_2_, …, *v*_|𝒱|−|𝒯 |_}, and terminal nodes 𝒯 = {*v*_|𝒱|−|𝒯 |+1_, …, *v*_|𝒱|_}. Likewise, **q** and **r** are separated into quantities defined on the root branch *e*_1_, the internal branches ℰ_int_ = {*e*_2_, …, *e*_|𝒱|−| 𝒯 |−1_} and terminal branches ℰ_erm_ = {*e*_|𝒱|−| 𝒯 |_, …, *e*_|𝒱|−1_}. In block form, the incidence and Laplacian operators are

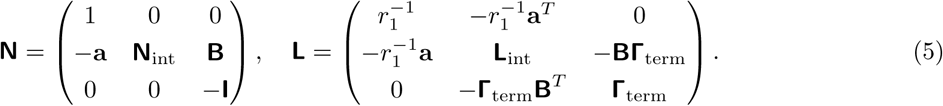

Here **a** = (1, 0, …, 0)^*T*^ and 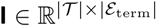 is the identity matrix. The rows of the matrices 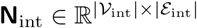 and 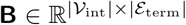 indicate the connections of the internal nodes *v*_*i*_ ∈𝒱_int_ to the internal branches *e*_*j*_ ∈ ℰ_int_ and the terminal branches *e*_*j*_ ∈ ℰ_term_ respectively. For brevity, we have denoted diag(**r**_term_)^−1^ *≡* **Γ**_term_. The internal Laplacian operator is given by 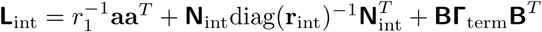.

The relation ∑_*i*_*N*_*ij*_ = 0 holds for all *j*, and so **N**^*T*^ **e**_|𝒱|_ = 0 where **e**_*M*_ *≡* (1, …, 1)^*T*^ ∈ ℝ^*M*^ denotes the all-one vector of size *M*, and so **e**_|𝒱|_ is the vector identifying all nodes. Thus, **e**_|𝒱|_ is a zero-eigenvalue mode of **L**, and multiplying (3) by 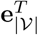 gives the condition for global mass conservation 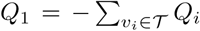. In fully connected networks, as considered here, this is the only zero-eigenvalue mode and so the conductance Laplacian **L** is rank |𝒱| − 1. It follows that |𝒯| + 1 boundary conditions are required to define the |𝒱| + |𝒯 | unknowns in (3) (|𝒱| node pressures and |𝒯 | terminal fluxes).

In the context of lungs we are generally interested in pressure boundary conditions on *P*_1_ and **P**_term_. To remove the arbitrary pressure constant, we first reformulate in terms of the pressure drop relative to the entry node *v*_1_ such that Δ**P** = *P*_1_**e**_|*V*|_ − **P**, and so Δ**P** = (0, Δ**P**_int_, Δ**P**_term_)^*T*^. Thus, given Δ**P**_term_ and inserting (4) into (3), the system of equations to be solved is

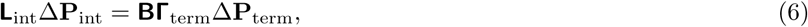

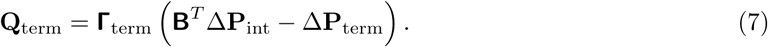

Equation (6) can be solved efficiently for Δ**P**_int_ by numerical linear algebra methods due to the sparsity of the operator **L**_int_. In general, for connected networks, **L**_int_ is invertible and so substituting (6) into (7) gives the terminal fluxes in terms of the pressure boundary conditions

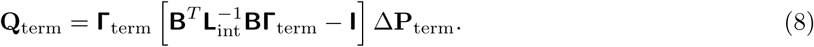

In practice, we solve this system of equations by implementing the “SparseLU” method from the C++ library Eigen [17], based on LU factorisation as detailed in section 1.1 of the Supplementary Text.

#### 2.1.1 Associated spectra of the Conductance Laplacian

The solution to (6) can also be expressed in terms of the eigenbasis of the operator **L**_int_. In general the internal Laplacian **L**_int_ has |𝒱| − |𝒯 | − 1 non-zero eigenvalues that we label *λ*_1_ ≤ *λ*_2_ ≤ … ≤ *λ*_|𝒱|−| 𝒯 |−1_ with corresponding eigenvectors **û**_1_, …, **û**_|𝒱|−| 𝒯 |−1_, where the hat indicates a normalised vector such that 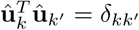 and *δ* is the Kronecker delta. Substituting the eigen-decomposition of **L**_int_ into (6) and solving for Δ**P**_int_ gives

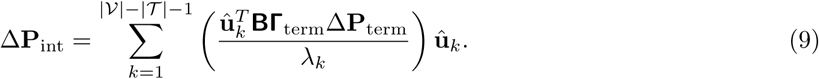

Substituting this solution into (7) gives the corresponding solution for **Q**_term_.

Dimensionality reduction can in principle be achieved using spectral methods by reconstructing the system with a subset of the eigenmodes. We can measure the relative dominance of each mode in the decomposition in (9) by the size of the coefficient 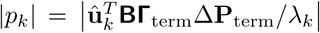. However, the relative weight of these modes is dependent on the boundary condition Δ**P**_term_. A more general measure of how well a subset of the eigenmodes captures the behaviour of the whole operator, independent of boundary conditions, is to consider how closely these modes approximate the inverse of the operator, **L**_int_, which we measure by the normalised distance *δ*_*L*_ [11] *given by*

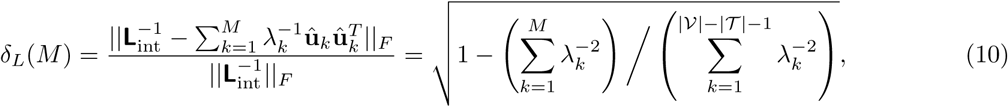

*where* ||.||_*F*_ indicates the Frobenius norm and *M ∈* {1, …, |𝒱| − |𝒯 | − 1} is the number of modes used in the approximation. Evidently, the modes with the smallest eigenvalues contribute the most to this reconstruction, however depending on the boundary conditions used they may not be the most dominant in the reconstruction of the solutions Δ**P**_int_ and **Q**_term_.

### 2.2 The Maury matrix

An alternative representation of the resistive airway tree network is presented in [15] for the case of a symmetric dyadic tree. Here, we generalise this operator to all connected trees and name it the Maury matrix in reference to [15]. To derive this operator, we begin by noting that connected sub-trees of the airway network can be defined due to the hierarchical structure of trees. We define the matrix **S** = (**s**_1_, …, **s**_|ℰ|_) where each column **s**_*j*_ maps the branch *e*_*j*_ to a subset of nodes 𝒱_*j*_ ⊂ {*v*_2_, …, *v*_|𝒱|_} descended from it such that

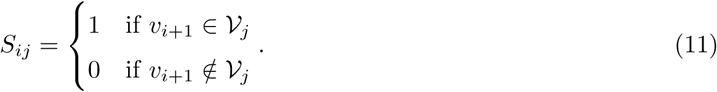

The first column of **S** is **s**_1_ = **e**_|𝒱|−1_ as all nodes except *v*_1_ are descended from the root branch. The column vectors **s**_*j*_ satisfy the relation 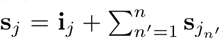,where 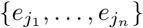 are the *n* child branches of *e*_*j*_, and **i**_*j*_ is the *j*th column of the identity matrix **I** ∈ ℝ^(| 𝒱|−1)*×*(| 𝒱|−1)^. This relation follows from noting that each sub-tree ℰ_*j*_ is a union of its child subtrees and its connecting node, such that 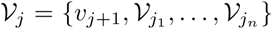. It follows that **SN**^*T*^ = (**e**_|𝒱|−1_, −**I**) and therefore

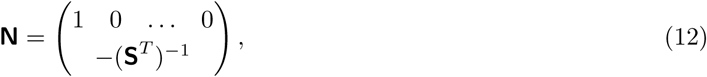

Substituting (12) into **Nq** = **Q** gives 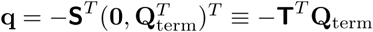, where the rows of **T** ∈ ℝ^|𝒯|*×*|ℰ|^ are comprised of the final |𝒯 | rows of the matrix **S**. Substituting this into (2) and multiplying both sides by −**S** diag(**r**) gives

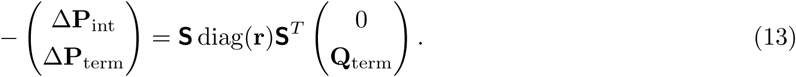

Comparing (13) to (3) we see that 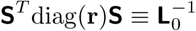 where 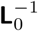 is the conductance Laplacian **L** with the first row and column removed (or the reduced Laplacian, as features in the matrix-tree theorem [18]). Finally, the bottom |𝒯| rows of (13) directly relate the terminal pressure drops to the fluxes

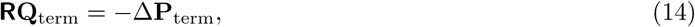

where **R** = **T**diag(**r**)**T**^*T*^ is the aforementioned Maury operator. Comparing with (8) gives

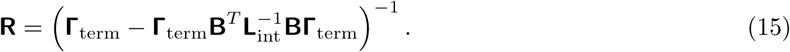

The entries of the terminal node map *T*_*ij*_ are 1 if the terminal node 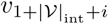 is descended from branch *e*_*j*_ and zero otherwise (see (11)). We denote the set of subset of terminal nodes descended from *e*_*j*_ as 𝒯_*j*_ ⊂ 𝒯 (see figure 1(a)). The Maury matrix is symmetric and in index notation its components are

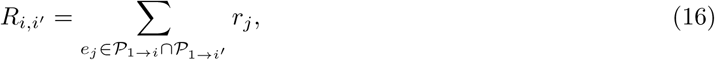

where 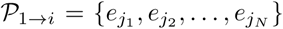 is the unique direct path on the tree from *v*_1_ to a terminal node *v*_*i*_ ∈𝒯. Each term 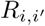 is equivalent to the resistance distance [19] between the node *v*_1_ and node 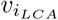 such that 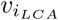 is the lowest common ancestor (LCA) node of the terminal nodes *v*_*i*_ and 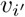.

A direct solve of (14) is an inefficient method for solving the system of equations (compared to (6) and (7)) because the matrix **R** has, in general, |𝒯 |^2^ non-zero entries and so cannot benefit from any algorithms optimised for the factorisation of sparse matrices. Nonetheless, this does not inhibit computation of the spectra of **R** because we can decompose **R** into sparse, non-square matrices. Details of the numerical methods used are given in section 1.2 of the Supplementary Text.

#### 2.2.1 Spectral properties of the Maury matrix

Diagonalising the Maury matrix **R** into its normalised eigenbasis and substituting into (14) gives

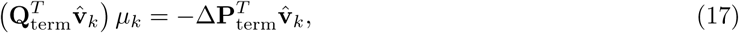

for each eigenvalue *µ*_*k*_ and associated eigenvector 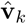. The matrix **R** has the form of an adjacency matrix for a complete network of resistors (with self-loops) relating the terminal nodes (as shown figure 1(b)). Therefore, as shown in (17), the modes act as resistors in parallel with resistance *µ*_*k*_, each subject to the pressure drop 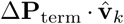. The matrix **R** is non-negative, regular and symmetric and so by the Perron–Frobenius theorem its largest eigenvalue mode *µ*_1_ is unique and the eigenvalues of the Maury matrix are all positive. Thus, we order the indices by *µ*_1_ ≥ *µ*_2_ ≥ … ≥ *µ*_|𝒯|_ (the opposite ordering to modes of the Laplacian).

From (17) the solution for the terminal node fluxes is

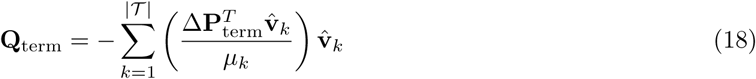

and so an approximation to 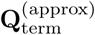 is given by limiting the sum in (18) to some subset of the mode spectrum. The magnitude of the coefficient 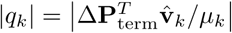 quantifies the relative importance of each term in the expansion of (18). As for the internal Laplacian operator, we measure the accuracy of such an approximation for arbitrary boundary conditions by the convergence of the inverse of **R** such that

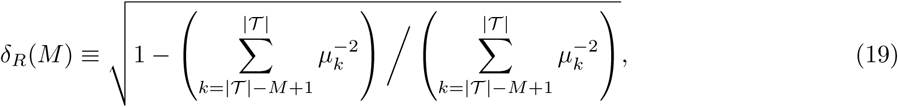

where *M* is the number of (smallest eigenvalue) modes used in the expansion. We will compare this with *δ*_*L*_(*M*) for specific networks to compare the general efficiency of dimensionality reduction for these two operators.

### 2.3 Dimensionality reduction in simulations of ventilation in the lungs

In models of ventilation in the lungs, the effects of lung tissue compliance generally dominate the mechanics compared to airway resistance, except in cases where there is airway narrowing or blockage [20]. Consider the simplest model of ventilation where each terminal node *v*_*i*_ ∈𝒯 is connected to an elastic bag of volume *V*_*i*_(*t*) and elastance *κ* with all bags subject to the same pleural pressure *P*_pl_(*t*) driving the breathing motion at time *t*. The terminal fluxes are related to the bag volumes via 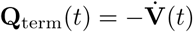 and the terminal pressures are **P**_term_ = [*P*_0_ + *P*_pl_(*t*)]**e**_|𝒯|_ + *κ***V**(*t*). Then (14) becomes

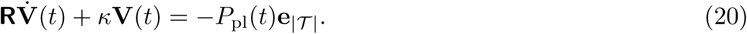

We are interested in characterising how the distribution of ventilation is affected by the airway resistance, and so have assumed that the lung unit compliances are all equal to *κ*. Diagonalising **R** into its eigenbasis in (20) gives

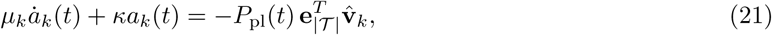

where 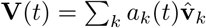. Thus each mode of **R** is characterised by an independent ODE with a resistance term *µ*_*k*_ and elastance *κ* driven by a pressure term proportional to 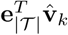. Solving (21) for *a*_*k*_(*t*) gives

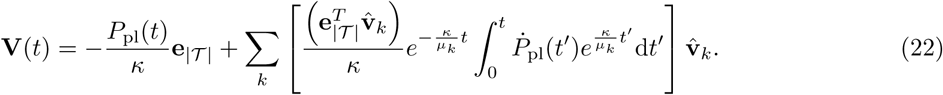

The first term in the solution is the elastic contribution from all units, while the second term is the resistive contribution. The sum in the resistive term can be approximated by the largest eigenvalue modes with a cutoff on the order *µ*_*k*_*/*(*κτ*) ∼ *O*(1) (where *τ* is the typical breath timescale), since modes satisfying *µ*_*k*_*/*(*κτ*) ≪ 1, will be dominated by lung unit compliance (the first term in (22)).

In the simulations used in this paper we take the pleural pressure to be sinusoidal such that *P*_pl_ = *P*_pl0_ + *P*_s_ sin(2*πt/τ*). Substituting into (22) we see that the periodic solution (when *t* ≫ *µ*_1_*/κ*) is

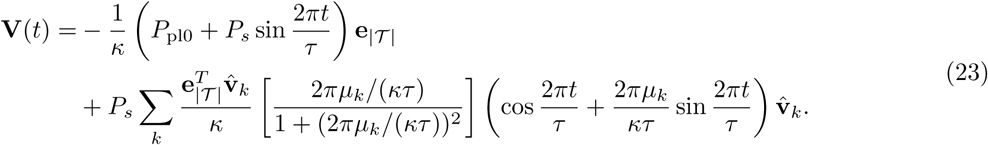

We define the ventilation of each terminal unit as Δ*V*_*i*_ = max (*V*_*i*_(*t*)) − min (*V*_*i*_(*t*)), where the maximum and minimum are taken over a single breath cycle 0 ≤ *t* < *τ*, which gives

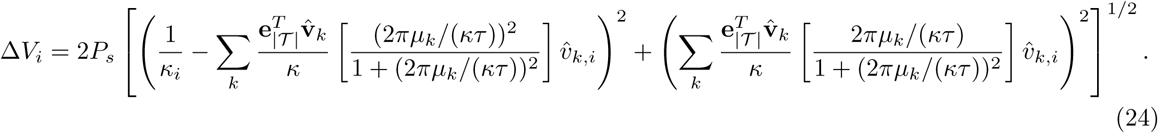

Inefficient ventilation of the lungs can occur if there are blockages or narrowing of the airway lumen, which can be caused by numerous pathophysiological factors, resulting in increased resistance of those airways and inhomogeneous delivery of gas to the lung units, commonly known as ventilation heterogeneity (VH). To characterise this in simulations we measure ventilation heterogeneity via the coefficient of variation of the ventilation *σ*_*V*_ across all terminal units,

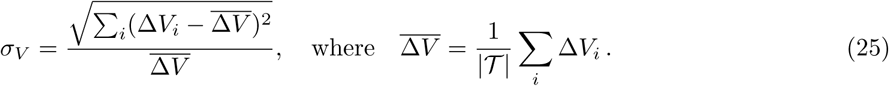

Note that the assumption of a sinusoidal pressure profile is used in this paper as it provides analytical solutions, however the above can be extended to generic pressure profiles simply by using numerical integration to estimate the value of the integral in (22). In many cases it would be desirable to set the pressure boundary condition to achieve a particular flow rate at the mouth. This results in a system of equations to be solved for the weights *a*_*k*_(*t*) and *P*_pl_(*t*), which again can be approximated by reducing to a subset of the largest eigenvalue modes.

Therefore the above decomposition gives a method to approximate the ventilation efficiency of an airway network given *κ* and *τ* by truncating the sums in (24) to *M* < |𝒯 | modes of the Maury matrix. While this method is specifically derived for sinusoidal breathing, it provides a useful qualitative measure of relative efficiency that can be used for comparing different networks and is easily computed using a subset of the full spectrum of the Maury operator.

### 2.4 Airway Models

In this paper we use idealised models of airway geometry and models derived from CT images, which we now briefly describe.

#### 2.4.1 Asymmetric Weibel branching

First, we adopt the Weibel-type model used in [21, 22] to systematically test the effect of asymmetry in resistance on the spectral properties of the Maury and Laplace operators. This idealised model describes a dyadic tree where each airway branches into major and minor daughters such that

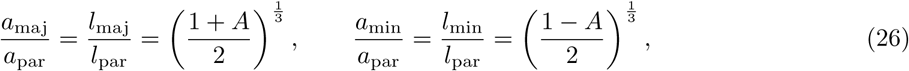

where *a* and *l* are the radii and lengths respectively of the parent and (major and minor) daughter branches as labelled by the subscripts. The parameter 0 ≤ *A* < 1 quantifies the asymmetry in branching (*A* = (1 − 2*r*) in terms of the notation in [21]). The cube root results in the two daughter airways having the same combined effective Poiseuille resistance and volume as the symmetric (*A* = 0) case. In this model we assume a length-to-diameter ratio of 3 for all airways and a fixed number of divisions *N* between the root branch and the terminal branches (resulting in *N* + 1 generations of airways).

#### 2.4.2 Horsfield model 1

In real lungs, branches terminate in a wide range of generations. To characterise this effect we use the Horsfield model 1 [23] derived from cast measurements of an adult male, with airways isotropically scaled to 60% of their original volume to a lung with ∼ 3L functional residual capacity (approximately that of the average adult). The model is terminated at Horsfield order 4 (as defined in [23]) because this corresponds to the transitional bronchioles as defined by Weibel [1] and gives 29, 240 terminal airways, approximately the number of acini in the lung [24]. The number of bifurcations from trachea to terminal bronchiole ranges from 10 to 25, this is achieved by imposing that parent airways branch into daughters with different ‘Horsfield orders’ (generations counting up from the terminal nodes). The radii and lengths are Gaussian random variables with mean taken from the data in [23] and a standard deviation of either 10 or 20% of the mean to approximate the variation in healthy lungs, which we label H10 and H20 respectively.

#### 2.4.3 Image-based (CT) models

Finally, we also use four image-based (IB) models derived from computed tomography (CT) images. These use volume-filling branching algorithms [25, 26] to populate the distal generations of airways in each lobe of the lung, which cannot be resolved from CT. The radius of these distal airways is assumed from an exponential fit to lung cast data, as in [26], such that

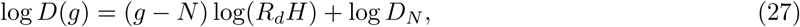

where *D*(*g*) is the airway diameter, *g* is the Horsfield order of the airway, *N* is the Horsfield order of the most distal ancestor airway taken from CT and its diameter *D*_*N*_. The parameter *R*_*d*_*H* is taken to be 1.15 as in [26].

The four IB models used here (labelled IB1-4) are from CT scans of children with cystic fibrosis and normal lung function (measured by spirometry) from a previously published study [27] (IB2 is shown in figures 4 and 5 below). The number of airways successfully acquired from CT depends on the image resolution, and was approximately between 3-5 generations of bifurcations from the trachea, and was acquired using the open-source software Pulmonary Toolkit [28]. The algorithm to generate these models used the C++ library `stl_reader` [29] and is detailed in section 1.3 of the Supplementary Text.

**Figure 2:**
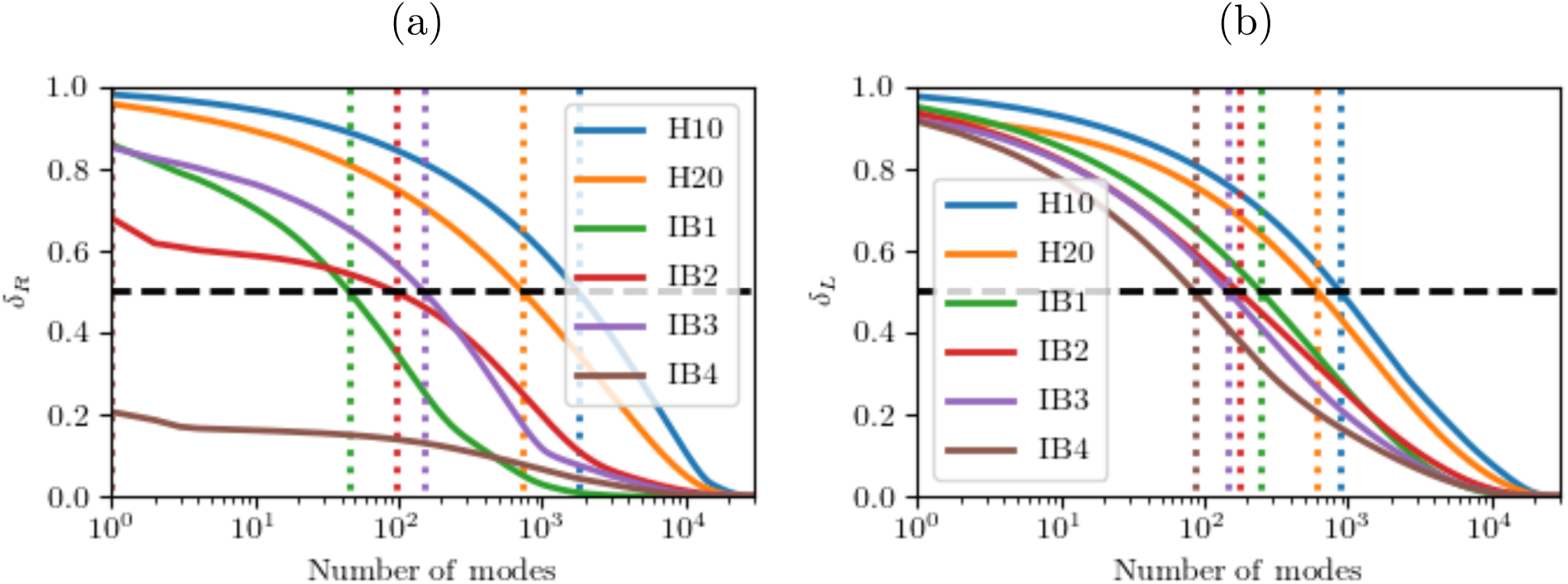
Normalised discrepancy in the cumulative estimate of the inverse of the (a) Maury operator and (b) internal Laplacian operator using the number of eigenmodes indicated on the *x*-axis where modes are ordered from smallest to largest eigenvalues. These are computed from the sums in (19) and (10) respectively. The networks H10 and H20 are the Horsfield model 1 with independent Gaussian random variables with standard deviations of 10% and 20% respectively added to the original airway diameter and length. Models IB1 through to IB4 are image-based trees derived from CT data as outlined in section 2.4. The vertical dashed lines indicate the point where each curve passes 50% accuracy.

**Figure 3:**
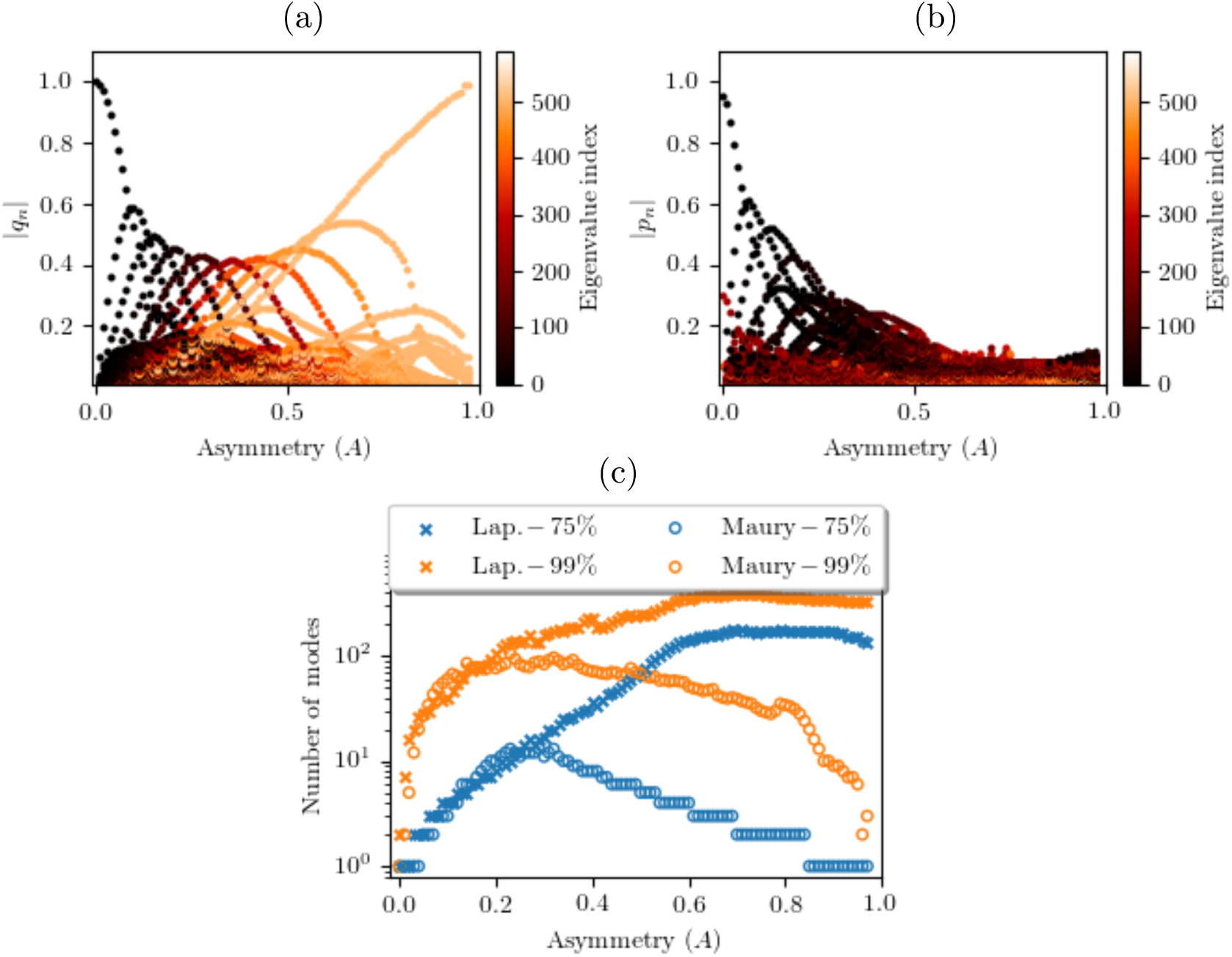
Relative magnitude of contributions of each eigenmode to the sums in equations (a) (18) and (b) (9) of a 10-generation Weibel network versus the *A* value of that network. Each dot represents the relative contribution of one of the 512 modes for the tree corresponding to that particular value of *A*, coloured by mode index; (a) Maury matrix (eigenvalues ordered largest to smallest) and (b) internal Laplacian (eigenvalues ordered smallest to largest). (c) The minimum number of modes required to achieve particular approximations the sums in (18) and (9) for the Maury and internal Laplace operators respectively.

**Figure 4:**
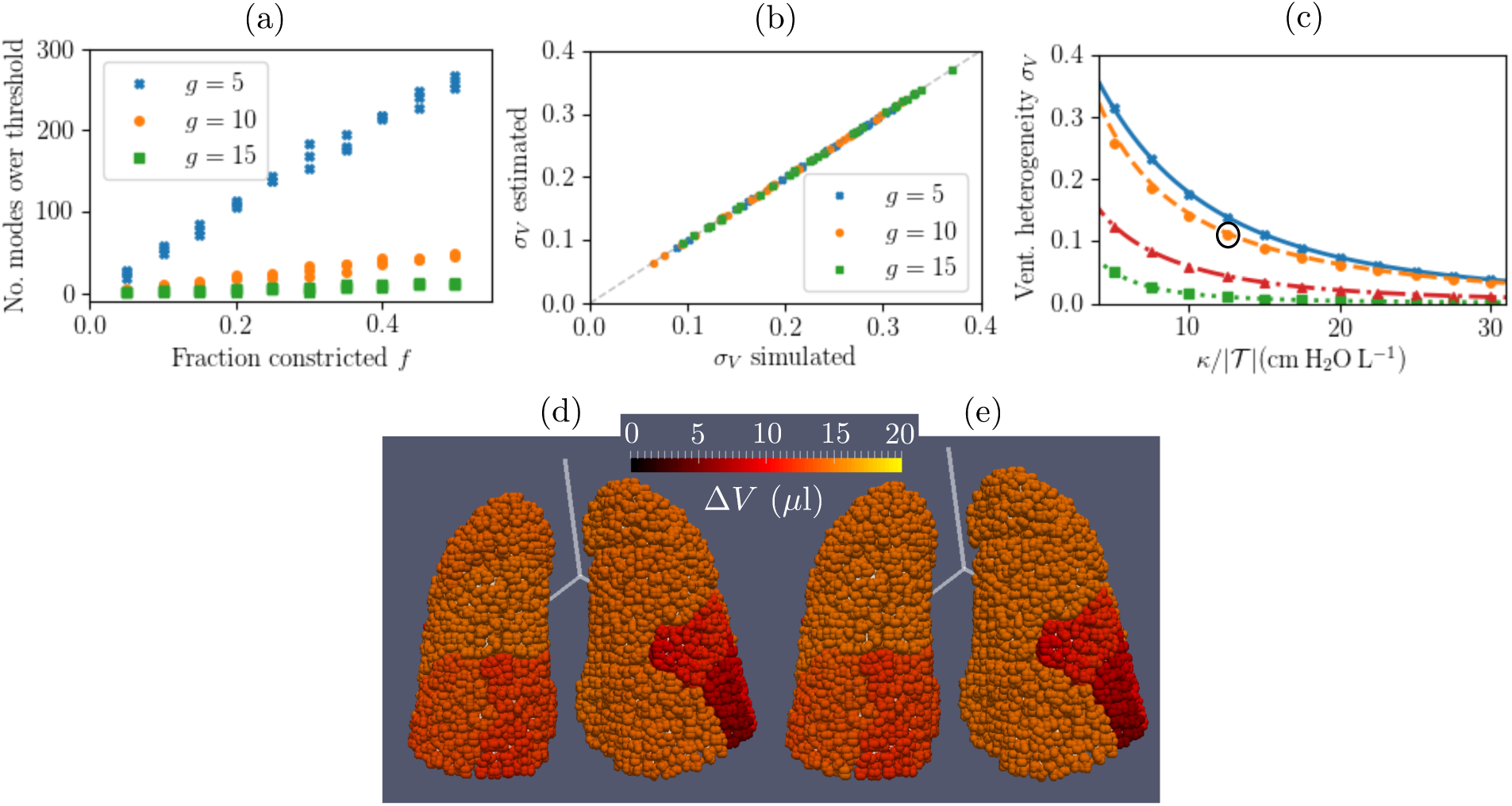
(a) Number of large eigenvalues (defined as *µ*_*k*_ *>* 0.1*κτ*) of the Maury matrix plotted against the fraction *f* of a given Horsfield order *g* constricted for the H10 airway tree network with added constrictions. The legend indicates which Horsfield order *g* constrictions were applied to for each tree. The total number of modes is 29,240. (b) Comparison of VH parameter *σ*_*V*_ for all networks in (a) computed through full simulation (*x*-axis) and estimated using just the largest-eigenvalue modes of the Maury matrix (*y*-axis) as identified in figure (a). All of the calculations in figures (a) and (b) used *κ/* |𝒯 | = 5cm H_2_0 L^−1^ and *τ* = 4s. (c) Simulated *σ*_*V*_ versus *κ/*|𝒯 | for the networks IB1-4 as labelled. Markers indicate results from numerical integration of (6) and (7) while lines indicate approximations using modes corresponding to the largest 1% of the eigenvalues in the Maury spectrum. (d) Visualisation of Δ**V** in the simulation of IB2 with *κ/*|𝒯 | = 12.5cmH_2_0L^−1^ (circled in (c)). (e) The prediction of Δ*V* for the same simulation using the 1% of the modes (corresponding to the largest 299 eigenvalues of the Maury matrix). See Supplementary Video 1 for a full 360*°*view of (d) and (e).

**Figure 5:**
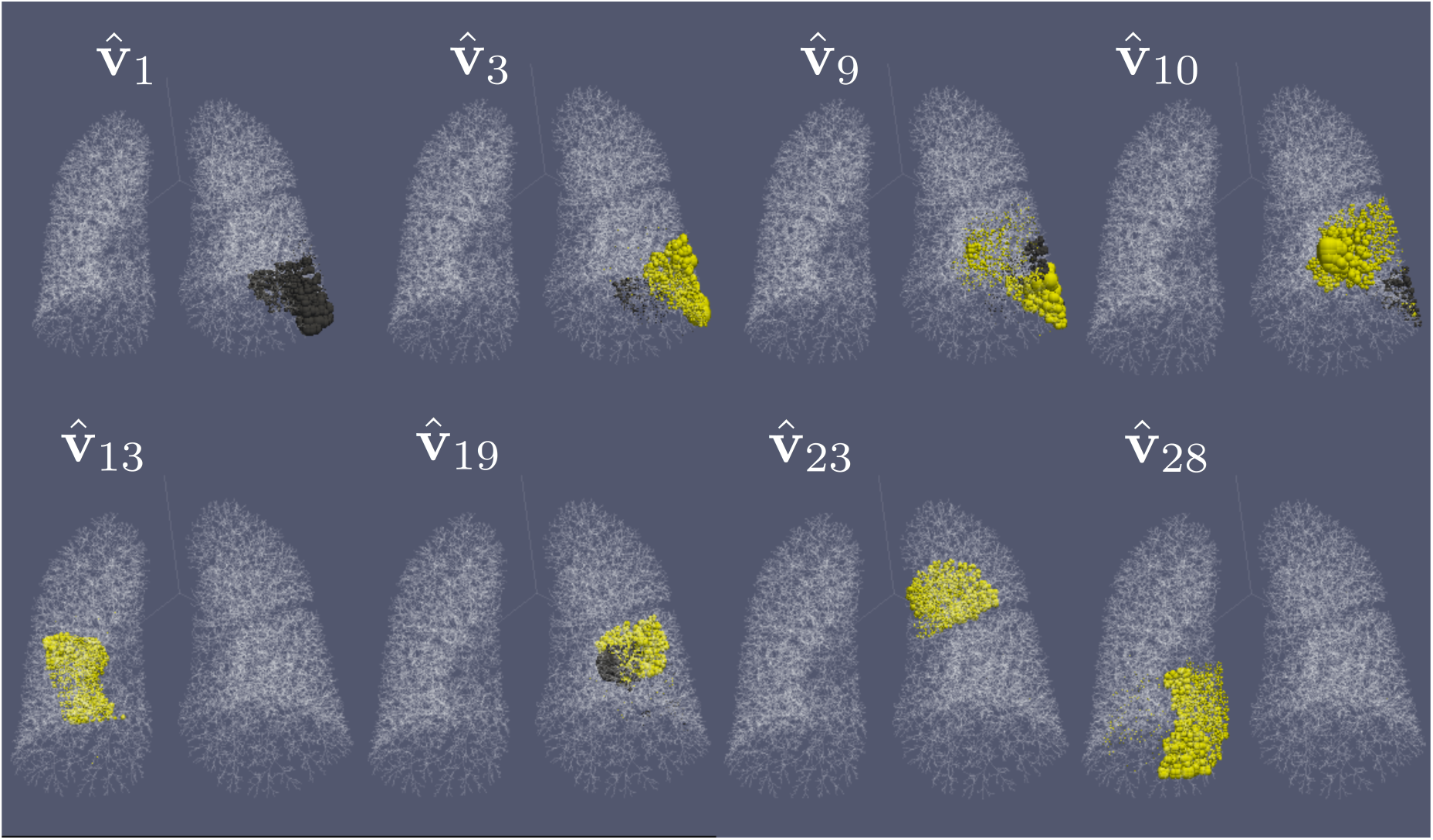
Visualisation of a selection of the eigenvectors corresponding to the largest eigenvalues of the Maury matrix for the IB2 network. The eigenvectors plotted are those with the largest values of 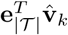 from the subset of modes that satisfy *µ*_*k*_ *>* 0.1*κτ* for the simulation values used in figure 4(d). In each plot, sphere size is proportional to the magnitude of eigenvector entries on all terminal nodes for the eigenvector labelled. The locations of the eigenvector entries highlight regions of impaired ventilation (according to the model) with the larger eigenvalue modes showing the least well-ventilated regions. The colours represent eigenvector entries with opposite sign. See Supplementary Video 2 for a full 360*°*view of this figure.

As a simple first approximation, the branch resistance is assumed to follow Poiseuille’s law and so *r* = 8*πvl/a*^4^ where *v* is the viscosity of air. To compute the spectra of the operators we use the Eigen [17] and Spectra [30] libraries in C++. The source codes of the algorithms used are available in online repositories [31, 32].

## 3 Results

In the following section we first study the spectra of the Maury operator for a number of idealised and image-based airway models and compare with the spectra of the internal Laplacian operator. Then, in the final results section we demonstrate how (24) can be used to produce a low-dimensional approximation of simulations and characterise realistic lung networks.

### 3.1 Comparison of Maury and internal Laplacian operator mode structures

Figure 2 shows the relative accuracy of the inverse approximation for each operator (from (19) and (10)) plotted against the number of eigenmodes used in the approximation. The Maury decomposition is more variable between networks, and in particular performs better (i.e. a smaller number of modes are required to capture more of the behaviour) for the image-based airway networks. In particular, the network labelled IB4 only requires a single mode to approximate 80% of the Maury inverse. This mode has a small eigenvalue and hence represents a “short-circuit” that, given appropriate boundary conditions, may account for the majority of the flow on the network. In contrast, the Frobenius norm of the inverse Laplace operator is largely comprised of high-resistance modes, and so the corresponding eigenvectors are centralised on high-resistance motifs in the network. The mode reconstruction is more accurate in the image-based networks than the Horsfield networks for either operator, which is due to the increased variance of resistance in these networks, however the effect is stronger for the Maury operator.

#### 3.1.1 Characterisation of network resistance with fixed pressure boundary conditions

We now consider fixed boundary conditions for the pressure-drop Δ*P*_term_ = **e**_|𝒯|_, such that flow is directed from the mouth to the alveolar region. Therefore, any flow heterogeneity that results is due to the distribution of resistance in the networks. To comprehensively cover a range of possible networks, we apply these conditions to the Weibel-type model in section 2.4.1 with *N* = 9 divisions (512 terminal nodes), representative of approximately 1*/*64th of the conducting airways [1].

As the network asymmetry parameter *A* is changed from *A* = 0 (symmetric) to *A* = 0.98 (all-but-one path having very large resistance), the dominant vectors of the Maury matrix (ranked by |*q*_*k*_|) transition from highest resistance modes (smallest index) to the lowest resistance modes (largest index), see figure 3(a). This characterises the transition from a flow solution that is homogeneously spread over all the terminal nodes and one that is concentrated on a small collection of terminal nodes that terminate the low-resistance paths in the tree. The *A* = 0 case is considered in more detail in section 1.4 of the Supplementary Text.

For all values of *A* there is a small subset of modes in the Maury spectrum that dominate the flow solution (figure 3(a)). In contrast, the Laplace spectrum does not demonstrate any strong ordering of mode dominance (ranked by |*p*_*k*_|) for the majority of *A* values and clear separation of modes occurs only at small values of *A* (figure 3(b)). For larger *A* values the Maury spectrum outperforms the Laplace spectrum in terms of the number of modes required to reconstruct the solutions (figure 3(c)). However, the difference is small in human lungs, where it is estimated that *A ≈* 0.35 [21]. The Maury matrix performs best at extreme values of *A*, but for all *A* values a 75% accuracy can be achieved using at most 10 eigenmodes (∼2% of the spectrum).

### 3.2 Practical application to simulations of lung ventilation mechanics

In this section we consider the simple mechanical model of regular tidal breathing introduced in section 2.3. In this case the largest eigenvalue modes of the Maury matrix dominate ventilation as they encode the high resistance paths in the tree (as demonstrated systematically in section 2.1 of the Supplementary Text).

#### 3.2.1 Dimensionality reduction in simulations and characterisation of realistic airway networks

First, we consider the Horsfield networks H10 and H20 (section 2.4.2), choosing parameters for elastance *κ* and time-scale *τ* that are representative of an average adult (listed in figure 4). In both cases, all eigenvalues of the Maury matrix satisfied *µ*_*k*_*/*(*κτ*) < 0.1. Therefore, the ventilation is approximately homogeneous in these networks and the airway resistance has little effect on ventilation.

To generate networks with high-resistance paths, we systematically applied constrictions to the Horsfield network by selecting a fraction *f* of the airways (at random) in a given Horsfield order *g* and reducing their radius by a factor between 50% and 95% (drawn from a uniform random distribution between these limits). The networks generated are idealised examples of the effects of obstructive lung disease, with different networks for disease centred on proximal (*g* = 15), central (*g* = 10), or distal (*g* = 5) airways. This is similar to constrictions applied in models of CF and asthma [33, 34, 35]. In the constricted networks, we found that as the number of random constrictions is increased (either by increasing *f* or reducing *g*), the number of independent large-eigenvalue modes of the Maury matrix rises (figure 4(a)). This agrees with our interpretation of these modes being centred on high-resistance paths in the tree.

The VH parameter *σ*_*V*_ from (25) does not depend strongly on *g* in this airway model, and so its variation in figure 4(b) is mainly due to the range of *f* values used (larger *f* results in more constricted airways and so greater VH). Regardless of *f* and *g* values, we see that using only large-eigenvalue modes in (25) gives a very good approximation of the simulated VH for all values of *f* and *g*. These reconstructions use at most 1% of the full Maury spectrum, and so represent a significant reduction in dimensionality.

Figure 4(c) shows the effect on VH of changing *κ* in image-based models (section 2.4.3) of ventilation and compares it to the prediction made using spectral decomposition. For each network we chose a range of realistic values of the total lung elastance *κ/* |𝒯 | for children from 5-30cm H_2_0 L^−1^ [36, 37, 38]. As *κ* is reduced, airway resistance has a more significant contribution to the dynamics in (20), and so VH is increased due to the heterogeneity of network resistance. The variation between the networks is characteristic of the different network structures, with IB1 and IB2 showing significantly more VH than IB3 and IB4.

The Maury spectral approximation (using 1% of the modes) in figure 4(c) performs marginally worse for smaller values of *κ*, however overall the agreement is very strong. The Maury eigenvalues and eigenvectors are unchanged by the parameter sweep in *κ* and so estimation of the ventilation distribution from equation (24) can be repeated for a range of *κ* values at negligible computational cost. As shown in table S1 (in Supplementary Text) the time to compute 1% of the eigenspectrum (as used here) is comparable to the time for a single simulation of the ventilation. Therefore, we were able to predict the relationship between VH and lung elastance accurately and more efficiently using this method.

An explicit example of this reduced dimensionality approximation is given in figures 4(d) and 4(e) for the network IB2 (corresponding to the highlighted simulation in 4(c)), which shows distributions of ventilation Δ*V*_*i*_ (see (24)) over terminal units computed by full simulation (figure 4(d)) and the reduced model (figure 4(e)). The heterogeneity in ventilation is a result of the airway geometry and topology generated by the space-filling branching algorithm, as well as the branch radii which scales with its parent airway from the CT image, and is captured very efficiently by the reduced model.

#### 3.2.2 Characterisation of realistic airway networks

Large-eigenvalue (high-resistance) eigenvectors of the Maury matrix are centralised on those terminal nodes most affected. Therefore, the eigen-decomposition of the Maury matrix is a powerful tool for visualising and ranking the least ventilated regions, regardless of the chosen value of lung elastance *κ*. Figure 5 visualises a subset of the largest eigenvalue eigenvectors of the IB2 network by the size of spheres superimposed onto its terminal nodes. Comparing this to figure 4(d) we see that these are localised on the least ventilated terminal nodes. Furthermore, the largest eigenvalue modes highlight the regions of poorest ventilation, as the eigenvalue is a measure of the resistance of the mode. The modes in figure 5 are selected by their value of 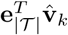, which is the coefficient in the mode sum in (24), so that they are all important in this reconstruction (for small enough *κτ*).

## 4 Discussion

In this paper we showed that in many realistic examples the Maury operator is a more efficient tool than the Laplacian operator for dimensionality reduction of the flow calculation on an airway network. Under fixed-pressure boundary conditions, it is not possible *a priori* to calculate only the eigenmodes which dominate the flow solution without computing the whole spectrum. However, when extended to simulations of lung ventilation, we found it is only the largest eigenvalue modes which significantly contribute to the extent of VH. These modes are generally the most efficient to compute using methods based on power iteration, as used here. Therefore, this provides a method for efficient prediction and characterisation of poorly ventilated regions of the lung based on the geometry and topology of the airway tree. The method has significant advantages over previous approaches of dimensionality reduction in airway models, which involve replacing parts of the tree with symmetric models [39, 40] based on “trumpet” models [41] of the airway geometry. These methods remove the asymmetry of the smaller airways, and sacrifice some of the complexity of the system to reduce dimensionality, which is not the case for the spectral graph theory methods used here.

The main limitation of this approach is the restriction to fixed linear resistance relations. This does not fully reflect the dynamics of ventilation in reality where airway resistance changes dynamically and non-linearly due to inertial effects [42]. Additionally, airways themselves are compliant [43], and dynamic bronchoconstriction occurs in lung conditions such as asthma changing the network resistance over time [44]. In cases where airway compliance becomes a dominant factor the Maury operator will cease to provide a useful description of this system. Nonetheless, the Maury matrix still has the same interpretation for a single instance of the resistance network, and so an interesting future study would be to study the dynamics of the Maury eigenspectrum in a system where airway resistance varies over time.

The results shown here are also limited to artificial lung airway geometries, using a combination of idealised cast-based models and algorithm-generated image-based networks. Therefore, the simulations and results in this paper do not predict how lungs will behave in specific states of health or disease. The models used do not contain enough information about the small airways or tissue compliance to make patient-specific predictions of VH, which was measured in the original study by ^3^He MRI [27]. Spectral methods, such as dynamic mode decomposition, have proven useful in similar contexts to generate low-dimensional characterisations of fluid dynamics simulations for machine learning image analysis applications [45]. In a similar way, the method here produces a low-dimensional representation of the airway tree resistance, which we aim to utilise in inverse modelling methods. For example, in future work we plan to fit CT-based models to ^3^He ventilation MRI data to develop patient-specific models of lung structure and function (similar to the approach in [34]), and this research will make it more feasible by reducing the computational time required to estimate the ventilation distribution for a candidate airway network over a range of parameters.

A further application of these findings is in the characterisation of complex airway tree models, such as those acquired from micro-CT imaging [46]. Micro-CT can only be performed on excised lungs, so it would not be possible to directly relate these to function. However, it will be interesting to characterise airway networks associated with various lung diseases. This work provides an efficient method for this as “problematic” lung regions can be visualised using the large-eigenvalue eigenmodes of the Maury matrix only.

The methods in this paper are generic to resistance trees, and hence we expect them to have applications outside of lung mechanics. First, other resistance trees in biology, such as the pulmonary vasculature, can be analysed using the same principles. Second, in discrete mathematics, the steady diffusion problem is identical to the linear resistance problem, therefore diffusion on tree networks can also be studied within this framework. For example, transport of gas in the respiratory zone of the lungs is dominated by diffusion on the tree network of acinar ducts, and so in future work we aim to use this formulation to produce low dimensional representations of models of these networks.

In summary, this paper has demonstrated how spectral graph theory provides a powerful tool for dimensionality reduction in the analysis of lung ventilation. This is achieved by reformulating the physical model for gas transport using a linear operator (the Maury matrix) that captures patterns of ventilation heterogeneity more efficiently than the traditional Laplacian operator, by exploiting the tree-like structure of the network. This approach shows potential for the description of a wider class of transport processes on tree-like networks.

## Acknowledgements

CW was supported by the UK Medical Research Council (MR/ R024944/1) and PL was funded by a University of Manchester internship. Imaging data was collected by the POLARIS group at University of Sheffield and was funded by the UK Medical Research Council (MR/M008894/1).

## References

[1] Weibel ER. 1963 Morphometry of the Human Lung. Springer-Verlag Berlin Heidelberg.

[2] Swan AJ, Clark AR, Tawhai MH. 2012 A computational model of the topographic distribution of ventilation in healthy human lungs. J. Theor. Biol. 300, 222–23. (doi: 10.1016/j.jtbi.2012.01.042)

[3] Kim M, Bordas R, Vos W, Hartley RA, Brightling CE, Kay D, Grau V, Burrowes KS. 2015 Dynamic flow characteristics in normal and asthmatic lungs. Int. J. Numer. Meth. Bio. 31, e02731. (doi: 10.1002/cnm.2730)

[4] Roth CJ, Ismail M, Yoshihara L, Wall WA. 2016 A comprehensive computational human lung model incorporating inter-acinar dependencies: Application to spontaneous breathing and mechanical ventilation. Int. J. Numer. Meth. Bio. 33, e02787. (doi: 10.1002/cnm.2787)

[5] Downie SR, Salome CM, Verbanck S, Thompson B, Berend N, King GG. 2007 Ventilation heterogeneity is a major determinant of airway hyperresponsiveness in asthma, independent of airway inflammation. Thorax 62, 684–689. (doi: 10.1136/thx.2006.069682)

[6] Horsley A, Wild JM. 2015 Ventilation heterogeneity and the benefits and challenges of multiple breath washout testing in patients with cystic fibrosis. Paediatr. Respir. Rev. 16, 15–18. (doi: 10.1016/j.prrv.2015.07.010)

[7] Farrow CE, Salome CM, Harris BE, Bailey DL, Berend N, King GG. 2017 Peripheral ventilation heterogeneity determines the extent of bronchoconstriction in asthma. J. Appl. Physiol. 123, 1188–1194. (doi: 10.1152/japplphysiol.00640.2016)

[8] Katifori E, Magnasco MO. 2012 Quantifying Loopy Network Architectures. PLoS ONE 7, e37994. (doi: 10.1371/journal.pone.0037994)

[9] Lee SH, Fricker MD, Porter MA. 2016 Mesoscale analyses of fungal networks as an approach for quantifying phenotypic traits. J. Complex Netw. 5, 145–159. (doi: 10.1093/comnet/cnv034)

[10] Meigel FJ, Alim K. 2018 Flow rate of transport network controls uniform metabolite supply to tissue. J. R. Soc. Interface 15, 20180075. (doi: 10.1098/rsif.2018.0075)

[11] Spielman DA. 2007 Spectral Graph Theory and its Applications. In 48th Annual IEEE Symposium on Foundations of Computer Science (FOCS07), IEEE. (doi: 10.1109/focs.2007.56)

[12] Forrow A, Woodhouse FG, Dunkel J. 2018 Functional Control of Network Dynamics Using Designed Laplacian Spectra. Physical Review X 8, 041043. (doi: 10.1103/physrevx.8.041043)

[13] Johnson OK, Lund JM, Critchfield TR. 2018 Spectral graph theory for characterization and homogenization of grain boundary networks. Acta Mater. 146, 42–54. (doi: 10.1016/j.actamat.2017.11.054)

[14] Brouwer AE, Haemers WH. 2011 Spectra of Graphs Springer-Verlag Berlin Heidelberg.

[15] Maury B. 2013 The Respiratory System in Equations. Springer Milan, Milano.

[16] Grady LJ, Polimeni JR. 2010 Discrete Calculus. Springer London. (doi: 10.1007/978-1-84996-290-2)

[17] Guennebaud G, Jacob B, Others. 2010 Eigen v3. Link: http://eigen.tuxfamily.org.

[18] Tutte WT. 2001 Graph Theory Cambridge : Cambridge University Press.

[19] Klein DJ, Randìc M. 1993 Resistance distance. J. Math. Chem. 12, 81–95. (doi: 10.1007/bf01164627)

[20] Kang W, Clark AR, Tawhai MH. 2018 Gravity outweighs the contribution of structure to passive ventilation-perfusion matching in the supine adult human lung. J. Appl. Physiol. 124, 23–33. (doi: 10.1152/japplphysiol.00791.2016)

[21] Majumdar A, Alencar AM, Buldyrev SV, Hantos Z, Lutchen KR, Stanley HE, Suki B. 2005 Relating Airway Diameter Distributions to Regular Branching Asymmetry in the Lung. Phys. Rev. Lett. 95, 168101. (doi: 10.1103/physrevlett.95.168101)

[22] Henry FS, Llapur CJ, Tsuda A, Tepper RS. 2012 Numerical Modelling and Analysis of Peripheral Airway Asymmetry and Ventilation in the Human Adult Lung. J. Biomech. Eng. 134, 061001. (doi: 10.1115/1.4006809)

[23] Horsfield K, Dart G, Olson DE, Filley GF, Cumming G. 1971 Models of the human bronchial tree. J. Appl. Physiol. 31, 207–217. (doi: 10.1152/jappl.1971.31.2.207)

[24] Haefeli-Bleuer B, Weibel ER. 1988 Morphometry of the human pulmonary acinus. Anat. Record 220, 401–414. (doi: 10.1002/ar.1092200410)

[25] Tawhai MH, Pullan AJ, Hunter PJ. 2000 Generation of an Anatomically Based Three-Dimensional Model of the Conducting Airways. Ann. Biomed. Eng. 28, 793–802. (doi: 10.1114/1.1289457)

[26] Bordas R, Lefevre C, Veeckmans B, Pitt-Francis J, Fetita C, Brightling CE, Kay D, Siddiqui S, Burrowes KS. 2015 Development and Analysis of Patient-Based Complete Conducting Airways Models. PLOS ONE 10, e0144105. (doi: 10.1371/journal.pone.0144105)

[27] Marshall H et al. 2017 Detection of early subclinical lung disease in children with cystic fibrosis by lung ventilation imaging with hyperpolarised gas MRI. Thorax 72, 760–762. (doi: 10.1136/thoraxjnl-2016-208948)

[28] Doel T. 2018. Pulmonary Toolkit v1.0. Link: https://github.com/tomdoel/pulmonarytoolkit.

[29] Reiter S. 2018. stl reader. Link: https://github.com/sreiter/stlreader.

[30] Qiu Y. 2015 Spectra. Link: https://spectralib.org/.

[31] Whitfield CA. 2020 Spectral properties of airway networks v1.0.2, 2020. Link: https://github.com/CarlWhitfield/Spectraldecomposition (doi: 10.5281/zenodo.3708114).

[32] Whitfield CA. 2020 Space filling airway branching algorithm v1.0.1, 2020. Link: https://github.com/CarlWhitfield/Space_filling_tree (doi: 10.5281/zenodo.3709073).

[33] Mitchell JH, Hoffman EA, Tawhai MH. 2012 Relating indices of inert gas washout to localised bronchoconstriction. Resp. Physiol. Neurobi. 183, 224–233. doi: 10.1016/j.resp.2012.06.031)

[34] Leary D et al. 2016 Hyperpolarized ^3^He magnetic resonance imaging ventilation defects in asthma: relationship to airway mechanics Physiol. Rep. 4(7), e12761 (doi: 10.14814/phy2.12761)

[35] Foy BH, Kay D, Borgas R. 2017 Modelling responses of the inert-gas washout and MRI to bronchoconstriction. Resp. Physiol. Neurobi. 235, 8–17. doi: 10.1016/j.resp.2016.09.009)

[36] Cook CD, Helliesen PJ, Agathon S. 1958 Relation Between Mechanics of Respiration, Lung Size and Body Size From Birth to Young Adulthood. J. Appl. Physiol. 13, 349–352. (doi: 10.1152/jappl.1958.13.3.349)

[37] Sharp JT, Druz WS, Balagot RC, Bandelin VR, Danon J. 1970 Total respiratory compliance in infants and children. J. Appl. Physiol. 29, 775–779. (doi: 10.1152/jappl.1970.29.6.775)

[38] Zapletal A, Paul T, Samanek M. 1976 Pulmonary elasticity in children and adolescents. J. Appl. Physiol. 40, 953–961. (doi: 10.1152/jappl.1976.40.6.953)

[39] Whitfield CA, Horsley A, Jensen OE. 2018 Modelling structural determinants of ventilation heterogeneity: A perturbative approach. PLOS ONE 13, e0208049. (doi: 10.1371/journal.pone.0208049)

[40] Hasler D, Anagnostopoulou P, Nyilas S, Latzin P, Schittny J, Obrist D. 2019 A multi-scale model of gas transport in the lung to study heterogeneous lung ventilation during the multiple-breath washout test. PLOS Computational Biology 15, e1007079. (doi: 10.1371/journal.pcbi.1007079)

[41] Paiva M. 1973 Gas transport in the human lung. J. Appl. Physiol. 35, 401–410. (doi: 10.1152/jappl.1973.35.3.401)

[42] Pedley TJ, Schroter RC, Sudlow MF. 1970 Energy losses and pressure drop in models of human airways. Resp. Physiol. 9, 371–386. (doi: 10.1016/0034-5687(70)90093-9)

[43] Lambert RK, Wilson TA, Hyatt RE, Rodarte JR. 1982 A computational model for expiratory flow. J. Appl. Physiol. 52, 44–56. (doi: 10.1152/jappl.1982.52.1.44)

[44] Venegas JG et al. 2005 Self-organized patchiness in asthma as a prelude to catastrophic shifts. Nature 434, 777–782. (doi: 10.1038/nature03490)

[45] Xi J, Zhao W 2019 Correlating exhaled aerosol images to small airway obstructive diseases: A study with dynamic mode decomposition and machine learning. PLOS ONE 14(1): e0211413. (doi: 10.1371/journal.pone.0211413)

[46] Boon et al. 2016 Morphometric analysis of explant lungs in cystic fibrosis. Am. J. Respir. Crit. Care Med. 193(5), 516–526. (doi: 10.1164/rccm.201507-1281OC)

