## Supplementary Text for "Spectral graph theory efficiently characterises ventilation heterogeneity in lung airway networks"

July 3, 2020

### 1 Supplementary methods

#### 1.1 Simulation of periodic ventilation

As detailed in section 2.3 of the main text, to simulate ventilation we define the boundary conditions

$$\mathbf{Q}_{\text{term}} = -\dot{\mathbf{V}}(t), \tag{S1}$$

$$\Delta\mathbf{P}_{\text{term}}(t) = -\kappa\mathbf{V}(t) - P_{\text{pl}}(t)\mathbf{e}_{|\mathcal{T}|} \tag{S2}$$

All of the simulations presented in the text use a sinusoidal pleural pressure $P_{\text{pl}}(t) = P_{\text{pl0}} + P_{\text{pl}}^{(s)}\sin(2\pi t/\tau)$. To simulate the periodic case (when $t \gg \tau$) we write

$$\Delta\mathbf{P}_{\text{int}}(t) = \Delta\mathbf{P}_{\text{int}}^{(0)} + \Delta\mathbf{P}_{\text{int}}^{(s)}\sin(2\pi t/\tau) + \Delta\mathbf{P}_{\text{int}}^{(c)}\cos(2\pi t/\tau), \tag{S3}$$

$$\Delta\mathbf{P}_{\text{term}}(t) = \Delta\mathbf{P}_{\text{term}}^{(0)} + \Delta\mathbf{P}_{\text{term}}^{(s)}\sin(2\pi t/\tau) + \Delta\mathbf{P}_{\text{term}}^{(c)}\cos(2\pi t/\tau), \tag{S4}$$

$$\mathbf{q}(t) = \mathbf{q}^{(0)} + \mathbf{q}^{(s)}\sin(2\pi t/\tau) + \mathbf{q}^{(c)}\cos(2\pi t/\tau), \tag{S5}$$

$$\mathbf{V}(t) = \mathbf{V}^{(0)} + \mathbf{V}^{(s)}\sin(2\pi t/\tau) + \mathbf{V}^{(c)}\cos(2\pi t/\tau), \tag{S6}$$

To ensure no net flow over a periodic cycle, $\mathbf{q}^{(0)} = 0$ must be satisfied. It follows from equation 2 in the main text ($\mathbf{q} = \text{diag}(\mathbf{r})^{-1}\mathsf{N}^T\mathbf{P}$) that $\Delta\mathbf{P}_{\text{int}}^{(0)} = 0$ and $\Delta\mathbf{P}_{\text{term}}^{(0)} = 0$. Therefore, from (S2), we find

$$\mathbf{V}^{(0)} = -\frac{P_{\text{pl}}^{(0)}\mathbf{e}_{|\mathcal{T}|}}{\kappa}. \tag{S7}$$

We define $P_{\text{pl}}^{(0)} \equiv -\kappa\left(V_{\text{FRC}} + V_T/2\right)/|\mathcal{T}|$ such that $V_{\text{FRC}}$ is the minimum lung volume during tidal breathing (in the limit of zero airway resistance). We also define $P_{\text{pl}}^{(s)} = -\kappa V_T/\left(2\,|\mathcal{T}|\right)$ so that $V_T$ is the tidal volume (in the limit of zero airway resistance). Substituting equations (S3)-(S6) into equations (6) and (7) (in the main text) and applying the boundary condition (S1) gives

$$\mathbf{L}_{\text{int}}\Delta\mathbf{P}^{(s)}_{\text{int}} - \mathbf{B}\boldsymbol{\Gamma}_{\text{term}}\Delta\mathbf{P}^{(s)}_{\text{term}} = 0, \tag{S8}$$

$$\boldsymbol{\Gamma}_{\text{term}}\left(\Delta\mathbf{P}^{(s)}_{\text{term}} - \mathbf{B}^T\Delta\mathbf{P}^{(s)}_{\text{int}}\right) - (2\pi/\tau)\mathbf{V}^{(c)} = 0, \tag{S9}$$

$$\mathbf{L}_{\text{int}}\Delta\mathbf{P}^{(c)}_{\text{int}} - \mathbf{B}\boldsymbol{\Gamma}_{\text{term}}\Delta\mathbf{P}^{(c)}_{\text{term}} = 0, \tag{S10}$$

$$\boldsymbol{\Gamma}_{\text{term}}\left(\Delta\mathbf{P}^{(c)}_{\text{term}} - \mathbf{B}^T\Delta\mathbf{P}^{(c)}_{\text{int}}\right) + (2\pi/\tau)\mathbf{V}^{(s)} = 0. \tag{S11}$$

We solve this system of equations using SparseLU decomposition solver in the Eigen C++ libraries by representing these equations as a single matrix system of the form $\mathbf{A}\mathbf{x} = \mathbf{b}$ as follows

$$\begin{pmatrix} \mathbf{L}_{\text{int}} & -\mathbf{B}\boldsymbol{\Gamma}_{\text{term}} & 0 & 0 & 0 & 0 \\ -\boldsymbol{\Gamma}_{\text{term}}\mathbf{B}^T & \boldsymbol{\Gamma}_{\text{term}} & 0 & 0 & 0 & -\left(\frac{2\pi}{\tau}\right)\mathbf{I} \\ 0 & 0 & \mathbf{L}_{\text{int}} & -\mathbf{B}\boldsymbol{\Gamma}_{\text{term}} & 0 & 0 \\ 0 & 0 & -\boldsymbol{\Gamma}_{\text{term}}\mathbf{B}^T & \boldsymbol{\Gamma}_{\text{term}} & \left(\frac{2\pi}{\tau}\right)\mathbf{I} & 0 \\ 0 & \mathbf{I} & 0 & 0 & \kappa\mathbf{I} & 0 \\ 0 & 0 & 0 & \mathbf{I} & 0 & \kappa\mathbf{I} \end{pmatrix} \begin{pmatrix} \Delta\mathbf{P}^{(s)}_{\text{int}} \\ \Delta\mathbf{P}^{(s)}_{\text{term}} \\ \Delta\mathbf{P}^{(c)}_{\text{int}} \\ \Delta\mathbf{P}^{(c)}_{\text{term}} \\ \mathbf{V}^{(s)} \\ \mathbf{V}^{(c)} \end{pmatrix} = \begin{pmatrix} 0 \\ 0 \\ 0 \\ 0 \\ \kappa V_T \mathbf{e}^T_{|\mathcal{T}|}/\left(2\,|\mathcal{T}|\right) \\ 0 \end{pmatrix}, \tag{S12}$$

where the final two rows are the boundary conditions in (S2).

#### 1.2 Computation of spectra

The spectra of the idealised networks are computed using the Lanczos algorithm which requires only repeated multiplication of a vector by the operator in question. In general the matrix $\mathbf{R}$ has $|\mathcal{T}|^2$ non-zero entries and so it is computationally inefficient to perform such multiplications directly. Instead, the multiplication operation $\mathbf{y} = \mathbf{R}\mathbf{v}$ can be computed efficiently for an arbitrary vector $\mathbf{v}$ using the sparse decomposition $\mathbf{R} = \mathbf{T}\text{diag}(\mathbf{r})\mathbf{T}^T$. Then, performing the sparse multiplication operations $\mathbf{w} = \mathbf{T}^T\mathbf{v}$, and $\mathbf{y} = \mathbf{T}\text{diag}(\mathbf{r})\mathbf{w}$ completes the operation. The number of operations required for each multiplication roughly scales as $|\mathcal{E}|\,N$ where $N$ is the number of airway generations in the network.

The Horsfield and CT-based networks have $\sim 30,000$ terminal nodes so we compute the full spectrum using a shift-solve method (see Spectra documentation for implementation [10]). To do this efficiently, for the matrix $\mathbf{A} \in \{\mathbf{R}, \mathbf{L}_{\text{int}}\}$ we use the following algorithm.

1. Set $n = n_0$ and $m = 0$.

2. Compute the $n+1$ largest eigenvalue modes of $\mathbf{M}$ using the Lanczos algorithm. The eigenvalues will be $\lambda_1, \ldots, \lambda_{n+1}$ in descending size

3. If $\lambda_n \neq \lambda_{n+1}$ define $x = (\lambda_n + \lambda_{n+1})/2$ and discard the mode $\lambda_{n+1}$.

4. Otherwise, increment $n = \min(n + n_0 + 1, N_e)$, where $N_e$ is the total number of modes to be found, and return to step 1.

5. Increment $m = m + n$ and set $n = \min(n_0, N_e - m)$.

6. Compute the $n+1$ largest eigenvalue modes of $\mathbf{B} = (x\mathbf{I} - \mathbf{A})^{-1}$ and define the eigenvalues as $\nu_1, \ldots, \nu_{n+1}$ in descending order such that the next $n+1$ modes of $\mathbf{A}$ are $\lambda_{m+i} = x - \nu_i^{-1}$ for $i = 1, \ldots, n+1$.

7. If $\lambda_{m+n} \neq \lambda_{m+n+1}$ define $x = (\lambda_{m+n} + \lambda_{m+n+1})/2$ and discard the mode $\lambda_{m+n+1}$.

8. Otherwise, increment $n = n + n_0$ and return to step 6.

9. If $m < N_e$, return to step 5.

When considering the Maury matrix $\mathbf{R}$, step 5 in the above algorithm uses the operation $\mathbf{y} = (x\mathbf{I} - \mathbf{R})^{-1}\mathbf{v}$ for an arbitrary constant $x$ and vector $\mathbf{v}$, which is expensive to solve for if using $\mathbf{R}$ directly. However, we can emulate this operation efficiently by solving equations (6) and (7) in the main text for the case where the terminal edge resistances are modified such that $r_j \to r_j - x$ for all $\{e_j = (v_i, v_{i'}) \in \mathcal{E} \mid v_{i'} \in \mathcal{T}\}$ and the pressure-drop is given by $\mathbf{P}_{\text{ext}} - P_1\mathbf{e} = \mathbf{v}$. Then, the operation is equivalent to $\mathbf{y} = -\mathbf{Q}_{\text{ext}}$ for this solution.

#### 1.3 Generation of CT image-based airway models

The image-based models used in this paper are based on the algorithms developed in [5] and [6]. Airway centreline models were generated automatically from the CT data from the cohort in [7] using Pulmonary Toolkit (PTK) [8]. Registration of the lobes was achieved semi-automatically, by correcting the initial guess generated by PTK to coincide with lobe fissures, which were identifiable in all images. The lobe surfaces were outputted and stored in `.stl` files. The branching algorithm was written in C++ and used the `stl_reader` header library [9]. The algorithm is initialised with the following steps

1. A point density $\rho$ is calculated by dividing the total volume of the lobes by 30,000 (approximate number of acini).

2. A 3D grid of points is generated with spacing $\rho^{1/3}$, and the position of each point is randomly perturbed by a uniformly distributed random number between $-0.01\rho^{1/3}$ and $0.01\rho^{1/3}$. This perturbation avoids potential overlap with the `.stl` gridpoints, which can cause problems in determining which points are inside or outside a given surface, as well as adding a small random element to the generated tree.

3. Each point is determined 'inside' or 'outside' of each lobe by casting a line and counting the number of surfaces crossed. Points deemed outside the whole lung are discarded.

4. The terminal nodes of the CT centreline model are identified and assigned to each lobe using the same method.

Starting from the terminal nodes of the CT centreline tree, the space-filling branching algorithm proceeds as follows:

1. The grid points within a lobe are assigned to the nearest open terminal node of the tree.

2. Each point cloud is split by a plane through the mean position of the points in the cloud. The normal of the plane is the cross product of the direction of the terminating branch and the direction from the terminal node to the point cloud mean position. In some cases, these directions may be degenerate, in which case the parent branch of the terminal branch is used instead of the terminal branch direction.

3. New branches are generated along the direction from the terminal node towards the centre of mass of each cloud and terminated 40% of the way.

4. The branching angle of each branch with the parent branch is checked, and if it exceeds z the branch is rotated to have angle $\pi/3$.

5. If the branch terminates outside the lobe surface, the branch is iteratively shortened until this is not longer the case.

6. If the branch is $< 1$mm in length, or its point cloud has 1 point in it, the branch is terminal and its terminal node is classed as 'closed'. The nearest point in its point cloud is removed from reassignment in the next round.

7. If open terminal nodes remain, return to step 1.

Once the branching is complete, diameters are assigned to the generated airways using equation (27) from the text, so that

$$\log D(x) = (x - N)\log(R_d H) + \log D_N, \tag{S13}$$

where $D(x)$ is the airway diameter, $x$ is the Horsfield order of the airway, $N$ is the Horsfield order of the most distal ancestor airway taken from CT and its diameter $D_N$.

#### 1.4 Spectral properties of the Maury matrix on symmetric dyadic trees and sub-trees

Consider the case where a subset of the terminal nodes $\mathcal{T}_j \subset \mathcal{T}$ of the tree are indistinguishable by symmetry, *i.e.* all paths from their lowest common ancestor (LCA) edge $e_j$ are identical in resistance structure. By symmetry, $\mathcal{T}_j$ can be replaced by a single terminal node connected to $v_i$ (where $e_j = \{v_i, v_{i'}\}$) by a single edge with resistance equal to the effective resistance of the replaced sub-tree. In fact, approximating distal sub-trees as symmetric is a method of dimensionality reduction in these systems [1, 2] which has its origin in so-called "trumpet" models of the airways [3]. However, we find that this is a specific example of the more general method of dimensionality reduction proposed here based on the Maury spectrum.

Airway trees are almost always dyadic (each branching point being a bifurcation), so we explicitly consider that case here. In the dyadic case the number of terminal nodes in the subtree $\mathcal{S}_j = \{\mathcal{V}_j, \mathcal{E}_j\}$ is $|\mathcal{T}_j| = 2^N$ where $N+1$ is the number of generations of airways in the sub-tree, *i.e.* there are $N$ bifurcations between the edge $e_j$ and the terminal nodes $\mathcal{T}_j$. If we assume the terminal nodes are ordered such that all $v_i \in \mathcal{T} \setminus \mathcal{T}_j$ are listed first, and $v_i \in \mathcal{T}_j$ are last then $\mathsf{R}$ becomes

$$\mathsf{R} = \begin{pmatrix} \mathsf{A} & \mathbf{r}_a \mathbf{e}_{2^N}^T \\ \mathbf{e}_{2^N} \mathbf{r}_a^T & \mathsf{Y} \end{pmatrix}. \tag{S14}$$

The matrix $\mathsf{A} \in \mathbb{R}^{(|\mathcal{T}|-2^N)\times(|\mathcal{T}|-2^N)}$ only contains terms corresponding to resistance paths that terminate outside of the sub-tree nodes $\mathcal{V}_j$. The off diagonal blocks contain the resistance of paths shared between terminal nodes not in the sub-tree ($v_i \in \mathcal{T} \setminus \mathcal{T}_j$) and those that are ($v_i \in \mathcal{T}_j$). These are all the same for all terminal nodes in the symmetric subtree ($v_i \in \mathcal{T}_j$) as they share the same common ancestor and, by definition, are more distantly related to the other terminal nodes ($v_i \in \mathcal{T} \setminus \mathcal{T}_j$). Therefore, these blocks can be expressed as the outer product of a vector of resistances $\mathbf{r}_a$ with the all-one vector $\mathbf{e}_{2^N}$.

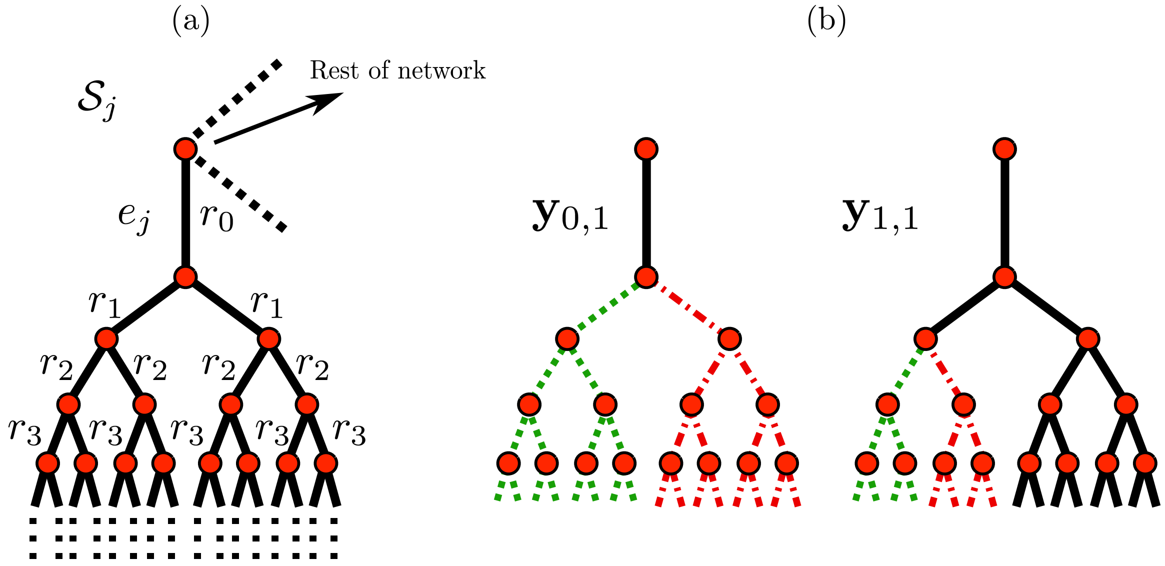


Figure S1: (a) Sketch of a symmetric sub-tree $\mathcal{S}_j$ with parent edge $e_j$ and equal resistance of all airways in each generation of the sub-tree ($n = \{0, 1, \ldots\}$. (b) Sketch of the eigenvectors $\mathbf{y}_{0,1}$ and $\mathbf{y}_{1,1}$ of the matrix $\mathbf{Y}$ corresponding to generations 0 and 1 of the sub-tree respectively. The green (dotted) and red (dot-dashed) lines indicate paths to terminal nodes whose corresponding entry in the eigenvector is $+1$ or $-1$ respectively (see equation (S16)).

The block $\mathbf{Y} \in \mathbb{R}^{2^N \times 2^N}$ contains resistance paths that terminate in the symmetric tree $\mathcal{S}_j$ and so has the structure

$$\mathbf{Y} = r_0 \mathbf{e}_{2^N} \mathbf{e}_{2^N}^T + r_1 \begin{pmatrix} \mathbf{e}_{2^{N-1}} \mathbf{e}_{2^{N-1}}^T & 0 \\ 0 & \mathbf{e}_{2^{N-1}} \mathbf{e}_{2^{N-1}}^T \end{pmatrix} + \ldots r_{N-1} \begin{pmatrix} \mathbf{e}_2 \mathbf{e}_2^T & 0 & \ldots & 0 \\ 0 & \ddots & \ddots & \vdots \\ \vdots & \ddots & \ddots & 0 \\ 0 & \ldots & 0 & \mathbf{e}_2 \mathbf{e}_2^T \end{pmatrix} + r_N \mathbf{I}, \tag{S15}$$

where $\mathbf{I} \in \mathbb{R}^{2^N \times 2^N}$ is the identity matrix. The resistances $r_n$ are the resistance of any given edge $e_j$ in generation $n \in \{0, \ldots, N\}$ of the subtree branches $\mathcal{E}_j$, as all the airway in a generation are identical by symmetry.

It follows from the definition above that the all-one vector $\mathbf{e}_{2^N}$ is an eigenvector of $\mathbf{Y}$. All other eigenvectors of $\mathbf{Y}$ can be given in (non-normalised) form as

$$\mathbf{y}_{g,m} = (\underbrace{0, \ldots, 0}_{(m-1)2^{N-g}}, \underbrace{1, \ldots, 1}_{2^{N-g-1}}, \underbrace{-1, \ldots, -1}_{2^{N-g-1}}, \underbrace{0, \ldots, 0}_{(2^g-m)2^{N-g}})^T \quad \forall\, g \in \{0, \ldots, N-1\},\ m \in \{1, \ldots, 2^g\}. \tag{S16}$$

The indices $g$ and $m$ are the airway generation and index respectively corresponding to the edge $e_{j'}$ in the subtree $\mathcal{S}_j$ as shown in figure S1(b). The vectors in (S16) satisfy $\mathbf{e}_{2^N} \cdot \mathbf{y}_{g,m} = 0$ and so, padded with zeros, are also eigenvectors of the matrix $\mathbf{R}$ in (S14) as shown below

$$\mathbf{R}\begin{pmatrix}\mathbf{0}\\ \mathbf{s}_{n,m}\end{pmatrix} = \begin{pmatrix}\mathbf{A0} + (\mathbf{e}_{2^N}^T\mathbf{y}_{g,m})\mathbf{r}_a\\ (\mathbf{r}_a^T\mathbf{0})\mathbf{e}_{2^N} + \mathbf{Y}\mathbf{y}_{g,m}\end{pmatrix} = \mu_g\begin{pmatrix}\mathbf{0}\\ \mathbf{y}_{g,m}\end{pmatrix}, \tag{S17}$$

where $\mu_g$ is the eigenvalue of $\mathbf{Y}$ corresponding to the eigenvector $\mathbf{y}_{g,m}$. These eigenvectors of $\mathbf{R}$ can also be expressed as

$$\hat{\mathbf{v}}_{g,m} = \frac{\mathbf{t}_{j_1} - \mathbf{t}_{j_2}}{|\mathbf{t}_{j_1} - \mathbf{t}_{j_2}|}, \tag{S18}$$

where $\mathbf{t}_{j_1}$ and $\mathbf{t}_{j_2}$ are the $j_1$th and $j_2$th columns of the matrix $\mathbf{T}$ (that indicate which terminal nodes are descended from the edges $e_{j_1}$ and $e_{j_2}$, the daughter edges of $e_{j'}$). Given symmetric boundary conditions on the symmetric sub-tree ($\Delta P_i$ = const for all $v_i \in \mathcal{T}_j$) these eigenvectors do not contribute to the flow solution at all (see equation (17) in the main text), reducing the effective dimensionality of this block from $2^N$ terminal nodes to 1, and so is equivalent to replacing the symmetric sub-tree with a single edge by Kirchhoff's law (*i.e.* a trumpet representation).

The above extends trivially to the case where an entire $N+1$ generation tree is symmetric, and so the whole tree is fully characterised by a single mode. As was shown in [4], the degeneracy of the $2^n$ modes associated with each generation $n = 0, \ldots, N-1$ means there are $N+1$ eigenvalues in total which can be labelled

$$\mu_{\text{symm,pf}} = 2^N R_{\text{eff}} \equiv \mu_1, \tag{S19}$$

$$\mu_{\text{symm},g} = 2^N\left(R_{\text{eff}} - R_{\text{trunc}}^{(g)}\right) \equiv \mu_k \;\forall\, k \in \{2^n+1, \ldots, 2^{n+1}\},\; n \in \{0, \ldots, N-1\}. \tag{S20}$$

where $R_{\text{trunc}}^{(g)} = \sum_{g'=0}^{g} r_{g'}/2^{g'}$ the effective resistance of the tree truncated at generation $g$ and $R_{\text{eff}} \equiv R_{\text{trunc}}^{(N)}$ is the effective resistance of the whole tree. The eigenvalue $\mu_{\text{symm,pf}} = \mu_1$ is the Perron–Frobenius eigenvalue and in the case of symmetric boundary conditions is the only one that contributes to the flux. All others again satisfy $\mathbf{e}\cdot\mathbf{v}_k = 0$. This also extends to symmetric $n$-adic trees as shown in the the following section, although we focus on the dyadic case in this paper.

To summarise, the spectrum of the Maury matrix, unlike the Laplacian spectrum, accounts for symmetry reductions exactly. However, these symmetric trees are an idealisation, and so dimensionality reduction by assuming symmetry is restricted to "nearly-symmetric" cases [1]. The spectral graph theory approach proposed here has no such restriction, and can be extended to arbitrarily asymmetric cases, as we show in the results section.

##### 1.4.1 Perturbation of the symmetric solution

Now consider the case where a single edge $e_j$ in generation $g_p$ of the otherwise symmetric dyadic tree is perturbed to have resistance $r_{g_p} + \delta r$. The Maury matrix is modified by the perturbation such that $\mathbf{R}_0 \to \mathbf{R}$ where

$$\mathbf{R} = \mathbf{R}_0 + \delta r\mathbf{t}_j\mathbf{t}_j^T. \tag{S21}$$

Multiplying by the eigenvector $\hat{\mathbf{v}}_{g,m}$ of matrix $\mathbf{R}_0$ (defined in the previous section) gives

$$\mathbf{R}\hat{\mathbf{v}}_{g,m} = \mu_{\text{symm},g}\hat{\mathbf{v}}_{g,m} + (\mathbf{t}_j^T\hat{\mathbf{v}}_{g,m})\delta r\,\mathbf{t}_j. \tag{S22}$$

Therefore, if $\mathbf{t}_j^T\hat{\mathbf{v}}_{g,m} = 0$ then $\hat{\mathbf{v}}_{g,m}$ is also an eigenvector of $\mathbf{R}$ and its eigenvector remains unchanged. First, this is satisfied for all of the eigenvectors where $g \geq g_p$ as these modes are by definition orthogonal to $\mathbf{t}_j$ (see (S18)). Second, this condition is also satisfied if the branch $e_j$ is not descended from the internal node associated with the mode $\hat{\mathbf{v}}_{g,m}$ (in which case, from (S18) all of the entries that are non-zero in $\mathbf{t}_j$ will be zero in $\hat{\mathbf{v}}_{g,m}$ and vice versa). Linearising the eigenvalue equation $\mathbf{R}\hat{\mathbf{v}}_k = \mu_k\hat{\mathbf{v}}_k$ for perturbations about the affected modes gives

$$(\mathbf{R}_0 - \mu_{\text{symm},g}\mathbf{I})\delta\mathbf{v}_{g,m} + (\delta r\mathbf{t}_j\mathbf{t}_j^T - \delta\mu_{g,m}\mathbf{I})\hat{\mathbf{v}}_{g,m} = 0, \tag{S23}$$

where we have used the notation $\hat{\mathbf{v}}_k = \hat{\mathbf{v}}_{g,m} + \delta\mathbf{v}_{g,m}$ and $\mu_k = \mu_{\text{symm},g} + \delta\mu_{g,m}$. The modes need to remain orthogonal to the unperturbed modes so the solutions for $\delta\mathbf{v}_{g,m}$ can be expressed as linear superpositions of the other symmetric modes $\hat{\mathbf{v}}_{g',m'}$. Furthermore, to retain orthonormality $\hat{\mathbf{v}}_{g,m}^T\delta\mathbf{v}_{g,m} = 0$. Therefore, we can express the eigenvector perturbations to linear order as

$$\delta\mathbf{v}_{g,m} \approx \sum_{g'=-1,g'\neq g}^{g_p-1} c_{g',m'}\hat{\mathbf{v}}_{g',m'}, \tag{S24}$$

where $m' = m(g')$ is the affected branch index for generation $g'$ and we have used the indices $g' = -1$ and $m' = 0$ to denote the Perron–Frobenius mode (which is not associated with an internal node). Substituting into (S23) and comparing coefficients we can solve for $c_{g',m'}$ to find

$$\delta\mathbf{v}_{g,m} \approx (\mathbf{t}_j^T\hat{\mathbf{v}}_{g,m})\delta r \sum_{g'=-1,g'\neq g}^{g_p-1} \left(\frac{\mathbf{t}_j^T\hat{\mathbf{v}}_{g',m'}}{\mu_g - \mu_{g'}}\right)\hat{\mathbf{v}}_{g',m'}, \tag{S25}$$

$$\delta\mu_{g,m} \approx (\mathbf{t}_j^T\hat{\mathbf{v}}_{g,m})^2\delta r\,. \tag{S26}$$

Equations (S25) and (S26) predict that the more distal eigenmodes (i.e. those associated with nodes fewer generations above the perturbation) will be most strongly perturbed in the linear limit (both their eigenvalue and eigenvectors).

### 2 Supplementary Results

#### 2.1 Eigenvector centralisation in the Maury matrix locates high-resistance paths in airway trees

As shown in detail in section 1.4, the Maury matrix has a number of special properties when part or all of the airway tree is symmetric. Considering fixed boundary conditions ($\Delta\mathbf{P}_{\text{term}} = \mathbf{e}_{|\mathcal{T}|}$) then when the entire tree is symmetric only one mode has a non-zero contribution ($q_k$) to the flow solution, and so the whole tree acts as a single resistor. If the resistance of a single edge in generation $g'$ of this symmetric tree is perturbed to break the symmetry of the whole network, the tree can still be divided into $g'+1$ symmetric sub-trees, which can each be represented as single resistors as shown in [1] and figure S2(a). In figure S2(b) we show

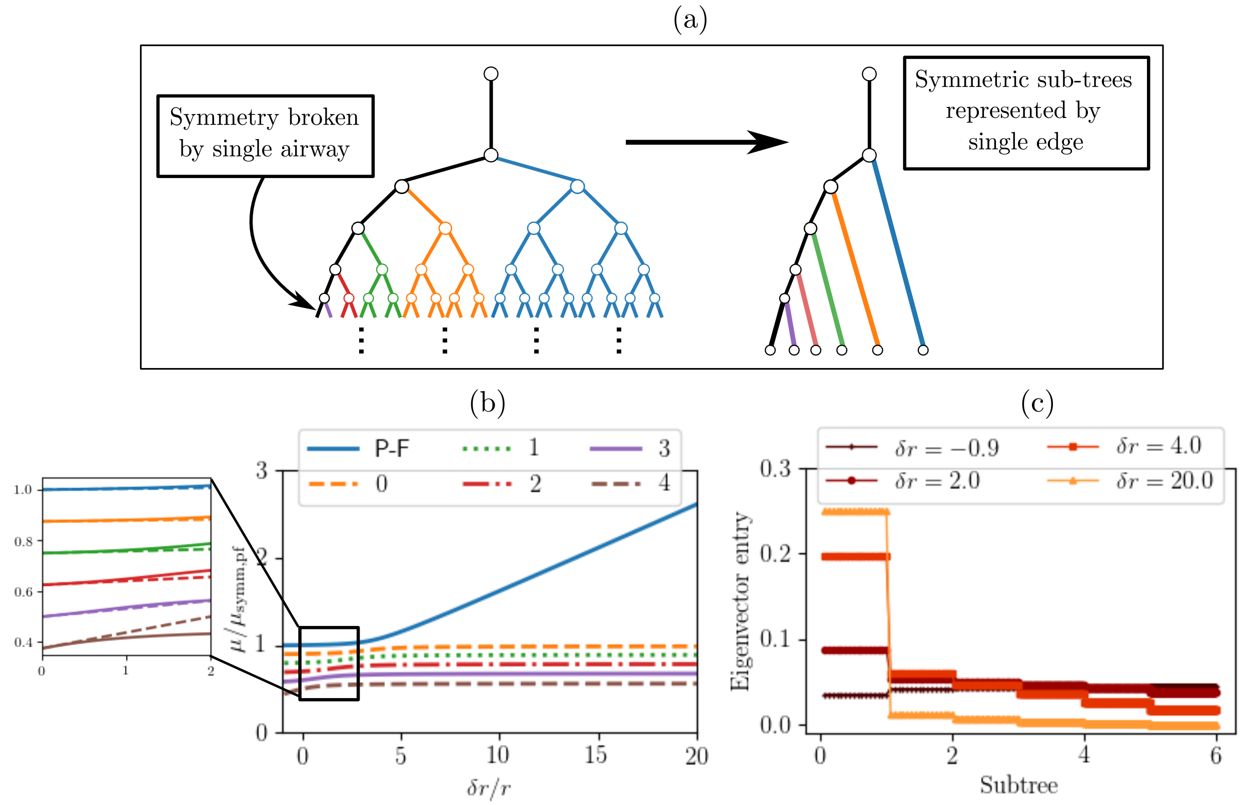


Figure S2: (a) Sketch of a network where all edges in each generation $g = 0, 1, \ldots$ have the same resistance except for the labelled edge in the 5th generation, which breaks the symmetry. Given symmetric boundary conditions on the terminal nodes, the flux solutions can be exactly reconstructed by replacing each symmetric subtree with a single resistor. (b) Response of the 6 affected eigenvalues of the Maury matrix for a symmetric tree with a resistance perturbation to an airway in generation 5 (as shown in (a)) of a 10-generation tree. The perturbation only affects the Perron-Frobenius mode $\mu_1$ (labelled P-F) and one mode for each generation $g \in \{0, \ldots, 4\}$ as labelled. Eigenvalues are normalised against the value of $\mu_{\text{symm,pf}}$ for the symmetric tree. The other 506 eigenmodes do not depend on $\delta r$ and, for symmetric boundary conditions, do not contribute to the flow solution. The inset shows the linear perturbation approximation in (S26) (dashed lines) against the simulated values (solid lines). (c) Entries of the eigenvector of the largest eigenvalue mode $\mu_1$ for various values of $\delta r$. The subtree numbers are defined as shown in figure (a) from left to right such that the terminal nodes downstream of the perturbation are plotted in the range (0,1), and those from the neighbouring sub-tree in the interval (1,2), and so on.

how the eigenvalues of the $g' + 1$ affected modes changes when the resistance of a single airway is perturbed. The initial trajectories of these curves can be predicted by linear perturbation theory in (S25) and (S26).

In figure S2(b) see that a large increase in resistance of a single airway is mostly captured by the largest eigenvalue mode and in figure S2(c) that the associated eigenvector becomes centralised on the nodes downstream of the affected airway. This corroborates with our interpretation of the largest eigenvalue modes of $\mathbf{R}$ indicating high resistance paths in the tree.

Thus large eigenvalue modes have eigenvectors centralised on high-resistance paths in the tree (when such paths exist) and small eigenvalue modes will be centralised on low-resistance paths (when they exist). However, only a small number of these modes will be important to the flux on the network depending on the boundary conditions. When the external pressure drop $\Delta\mathbf{P}_{\text{term}}$ is perpendicular to a mode ($\Delta\mathbf{P}_{\text{term}}^T \hat{\mathbf{v}}_k = 0$)

that mode will not contribute to the flow on the network.

#### 2.2 Matrix dimensions and computation times

Table S1 summarises some of the network sizes and simulation times used in the text for context. In the Horsfield networks (H10 and H20) we demonstrate the cases with no constrictions applied ($f = 0$), and hence the fewest large eigenvalue modes, and the case with the largest number of large eigenvalue modes in figure 4(a) of the main text.

| Network | Terminal nodes $\|\mathcal{T}\|$ | Sim. time (s) (Sinusoidal) | Computation time (s) | |
|---|---|---|---|---|
| | | | 1% eigenspectrum | $\mu_k > 2\,\|\mathcal{T}\|$ cmH$_2$OL$^{-1}$s |
| IB1 | 30050 | 20.9 ± 0.7 | 16.6 ± 0.3 | 39.2 ± 0.7 |
| IB2 | 29991 | 21.4 ± 0.2 | 15.9 ± 0.03 | 22.3 ± 0.4 |
| IB3 | 29893 | 21.0 ± 0.3 | 15.4 ± 0.3 | 0.424 ± 0.009 |
| IB4 | 29933 | 21.4 ± 0.1 | 15.9 ± 0.3 | 6.16 ± 0.04 |
| H10 ($f = 0$) | 29240 | 18.7 ± 0.2 | 14.7 ± 0.2 | 0.441 ± 0.010 |
| H20 ($f = 0$) | 29240 | 19.0 ± 0.3 | 14.7 ± 0.1 | 0.363 ± 0.006 |
| H10 ($f = 0.5$, $g = 5$) | 29240 | 18.8 ± 0.1 | 14.1 ± 0.3 | 12.7 ± 0.3 |

Table S1: Each simulation or computation time is the mean of 3 runs ± standard deviation on a single 2.6GHz processor. The simulations solve the system of equations in (S12). The last two columns show the time taken to compute the modes related to the largest 1% of eigenvalues of the Maury matrix, and to compute the modes above the cut-off value stated (corresponding to the minimum value of $\kappa\tau$ used in the paper).

### References


[1] Whitfield CA, Horsley A, Jensen OE. 2018 Modelling structural determinants of ventilation heterogeneity: A perturbative approach. PLOS ONE 13, e0208049. (doi:10.1371/journal.pone.0208049)

[2] Hasler D, Anagnostopoulou P, Nyilas S, Latzin P, Schittny J, Obrist D. 2019 A multi-scale model of gas transport in the lung to study heterogeneous lung ventilation during the multiple-breath washout test. PLOS Computational Biology 15, e1007079. (doi: 10.1371/journal.pcbi.1007079)

[3] Paiva M. 1973 Gas transport in the human lung. J. Appl. Physiol. 35, 401–410. (doi:10.1152/jappl.1973.35.3.401)

[4] Maury B. 2013 The Respiratory System in Equations. Springer Milan, Milano.

[5] Tawhai MH, Pullan AJ, Hunter PJ. 2000 Generation of an Anatomically Based Three-Dimensional Model of the Conducting Airways. Ann. Biomed. Eng. 28, 793–802. (doi:10.1114/1.1289457)

[6] Bordas R, Lefevre C, Veeckmans B, Pitt-Francis J, Fetita C, Brightling CE, Kay D, Siddiqui S, Burrowes KS. 2015 Development and Analysis of Patient-Based Complete Conducting Airways Models. PLOS ONE 10, e0144105. (doi:10.1371/journal.pone.0144105)

[7] Marshall H et al. 2017 Detection of early subclinical lung disease in children with cystic fibrosis by lung ventilation imaging with hyperpolarised gas MRI. Thorax 72, 760–762. (doi:10.1136/thoraxjnl-2016-208948)

[8] Doel T. 2018. Pulmonary Toolkit v1.0. Link: https://github.com/tomdoel/pulmonarytoolkit .

[9] Reiter S. 2018. stl_reader. Link: https://github.com/sreiter/stl_reader.

[10] Qiu Y. 2015 Spectra. Link: https://spectralib.org/.
